# A Galbonolide-producing *Streptomyces* reconfigures the plant root microbiota by activating salicylate-dependent defence metabolism

**DOI:** 10.64898/2026.08.27.747519

**Authors:** Clément Nicolle, Mohamed Zouaoui, Rémi Pendaries, Aurélien Amiel, Quentin Bazerque, Guillaume Marti, Bernard Dumas, Thomas Rey

## Abstract

*Streptomyces* sp. AgN23 is an epiphytic rhizobacterium that establishes in the *Arabidopsis* rhizosphere by activating plant immune responses. This activity depends on the secretion of polyketide galbonolides, which inhibit host inositol phosphoceramide synthase (IPCS) and thereby perturb sphingolipid homeostasis. However, the downstream signalling events linking IPCS inhibition to AgN23 enrichment in the rhizosphere remain unclear. Here, we show that AgN23 activates ethylene- and salicylic acid-dependent immune signalling, leading to coordinated stimulation of phenylalanine- and tryptophan-derived secondary metabolism. Using *Arabidopsis* mutants defective in these pathways, we show that these metabolites mitigate AgN23-induced root growth inhibition. We further show that the *npr1* mutant is strongly compromised in AgN23-triggered secondary metabolic responses, resulting in reduced rhizosphere colonization by AgN23. By comparing rhizosphere microbiota from wild-type and *npr1* plants, we distinguished direct AgN23 effects linked to intermicrobial competition from indirect effects mediated by host metabolic activation. In particular, AgN23 colonization occurred at the expense of several *Streptomycetaceae* ASVs and coincided with changes in bacterial and fungal taxa belonging to *Flavobacteriaceae* and *Mucoromycota*. Together, these findings define a mechanistic framework in which *Streptomyces* AgN23 interacts with NPR1-dependent signalling to reprogram root metabolism and rhizosphere community structure, notably through the production of specialized metabolites such as galbonolides.

**HIGHLIGHTS:**

- AgN23 root activity requires functional salicylate and ethylene signalling, but not jasmonate signalling.
- Salicylate pathway activation supports AgN23 rhizosphere colonization.
- AgN23 induces phenylalanine- and tryptophan-derived secondary metabolites that alleviate root growth inhibition.
- AgN23 reprogramming of the rhizosphere microbiota largely relies on NPR1 mediated responses to the *Streptomyces*.

**SIGNIFICANCE:** Plant roots interact with diverse microbial communities that support nutrition and enhance resilience to abiotic and biotic stresses. Engineering these communities is increasingly viewed as a key strategy for sustainable crop production. Beneficial microbes are often recruited through root exudation in a plant-driven “cry for help” process, yet the molecular mechanisms guiding this selective assembly remain incompletely understood. Here, we show that *Streptomyces* AgN23, a beneficial soil bacterium that produces antimicrobial and plant defence elicitor compounds such as galbonolides, exploits host immune signalling to establish itself in the rhizosphere and remodel the surrounding microbiota. AgN23 activates ethylene- and salicylic acid-dependent pathways in *Arabidopsis*, requiring the key signalling components EIN2 and NPR1, respectively. This immune activation triggers coordinated reprogramming of root metabolism that facilitates AgN23 growth, while also promoting phenylalanine- and tryptophan-derived secondary metabolites that mitigate root growth inhibition triggered by AgN23. By manipulating host metabolic outputs, AgN23 influences the composition of the root-associated community and secures its ecological niche. Given that *Streptomyces* species are consistently enriched in plants exposed to pathogens or drought, our findings support a model in which stress-associated *Streptomyces* leverage plant signalling hubs to thrive in the rhizosphere and participate in the assembly of a protective microbiota.

Graphical abstract
Model summarizing how galbonolides secreted by *Streptomyces* AgN23 activate ethylene and salicylic acid signalling pathways in *Arabidopsis*, leading to a coordinated reprogramming of root metabolism that supports AgN23 development in the rhizosphere and reshapes the resident microbiota. Galbonolide-induced responses require the phytohormone signalling components EIN2 and NPR1. NPR1 is required to induce the biosynthesis and rhizosphere release of exudates that promote AgN23 growth. In parallel, *Streptomyces*-triggered production of phenylalanine- and tryptophan-derived secondary metabolites mitigates root growth inhibition and contributes to rhizosphere microbiota restructuring.

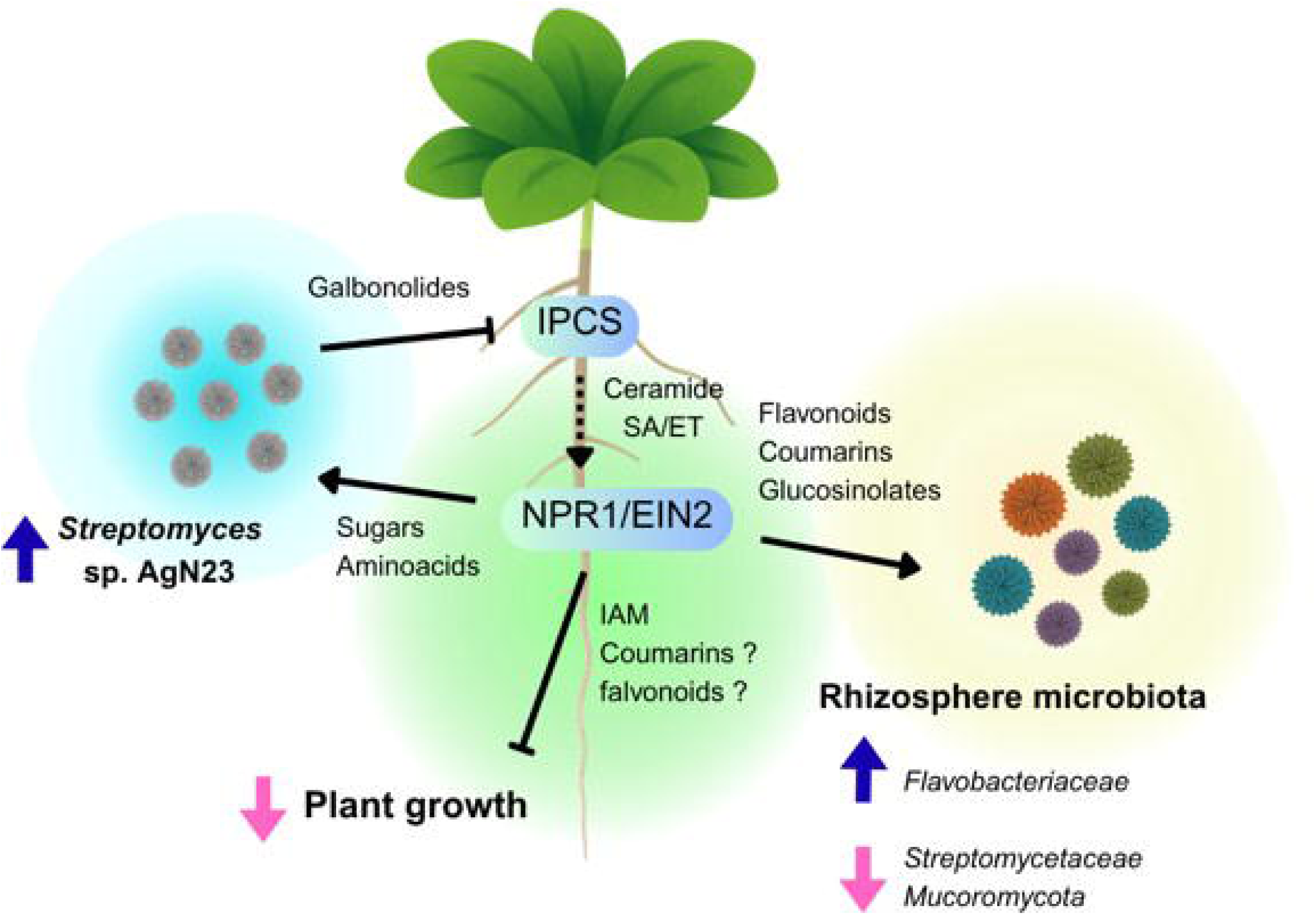

## INTRODUCTION

Plant roots are surrounded by abundant soil microbial communities, including bacteria, fungi, and oomycetes (Durán, Thiergart et al. 2018). This microbiota forms complex and dynamic assemblages that interact through mutualistic, commensal, and pathogenic relationships (Berg, Rybakova et al. 2020). Much like the gut microbiota in animals, the plant root microbiota contributes to host nutrition, development, and immunity (Getzke, Thiergart et al. 2019, Banerjee and van der Heijden 2023). Consequently, root-associated microbial communities are critical determinants of plant resilience to environmental stresses, and engineering the root microbiota holds great promise for sustaining agricultural productivity while reducing reliance on agrochemicals (Yu, Liu et al. 2022, Russ, Fitzpatrick et al. 2023).

Both bulk soil and rhizosphere microbiota are dominated by similar bacterial taxa, notably *Proteobacteria*, *Bacteroidetes*, *Firmicutes*, *Actinobacteria*, *Acidobacteria*, and *Verrucomicrobia* (Trivedi, Leach et al. 2020). However, their relative abundances differ between soil and the rhizosphere and fluctuate over plant development (Chaparro, Badri et al. 2014, Lu, Lu et al. 2025). Moreover, species within the same genus can exhibit markedly distinct traits depending on whether they inhabit bulk soil or are adapted to the rhizosphere niche (Ling, Wang et al. 2022). Recruitment of the rhizosphere microbiota is thought to result from interactions between plant-driven selection mechanisms and “hub” or “keystone” microorganisms (Agler, Ruhe et al. 2016). These organisms are part of the core root microbiota and can shape community structure through interactions with both the host plant and other microbial members. Their removal can lead to altered community assembly, commonly referred to as dysbiosis (Lee, Kong et al. 2021). Assembly of root and rhizosphere microbiota is also influenced by plant developmental processes and responses to biotic and abiotic stresses (Sharma, Kashyap et al. 2023). Plants can activate “cry for help” responses under drought or pathogen attack, promoting the recruitment of beneficial microorganisms that alleviate stress (Xu, Naylor et al. 2018, Stringlis, de Jonge et al. 2019, Stassen, Hsu et al. 2021). These microorganisms may produce phytohormones, modulate host signaling pathways, or synthesize antimicrobial compounds that contribute to plant defense (Carrión, Perez-Jaramillo et al. 2019, Finkel, Salas-González et al. 2020, Gu, Wei et al. 2020). Several bacterial genera, including *Streptomyces* spp., play key roles in this so-called plant extended immune system (Gao, Xiong et al. 2021, Pieterse 2025, Chen, Feng et al. 2026).

Although the molecular basis of stress-driven microbial recruitment remains incompletely understood, recent studies highlight the central role of root exudates in shaping microbiota composition (Yuan, Zhao et al. 2018, McLaughlin, Zhalnina et al. 2023). Among the diverse metabolites released by roots, certain compounds may act as “hub” signals exerting an extensive influence on microbial community structure and function (Brakhage 2024). For example, terpenes, phenylpropanoids, and indoles contribute to shaping rhizosphere microbiota by creating a distinctive chemical environment that differentiates it from bulk soil (Huang, Jiang et al. 2019). In addition, microbial interactions such as competition and cooperation further refine community composition by selectively excluding or recruiting specific strains (Kost, Patil et al. 2023). In *Arabidopsis*, root exudation includes methionine- and tryptophan-derived glucosinolates, as well as phenylalanine-derived flavonoids and coumarins (Voges, Bai et al. 2019). The coumarin scopoletin selectively inhibits fungal pathogens and certain bacterial taxa while promoting colonization by plant growth-promoting rhizobacteria (PGPR) and beneficial fungi (Berendsen, Vismans et al. 2018, Stringlis, Yu et al. 2018). In turn, colonization by these beneficial microbes activates salicylic acid, ethylene, and jasmonate signaling pathways, thereby enhancing plant immunity (Pieterse, Zamioudis et al. 2014, Mashabela, Piater et al. 2022).

*Streptomyces* spp. are dominant members of the Actinobacteria associated with plant roots and rhizospheres, representing approximately 10–30% of the total bacterial community (Bulgarelli, Rott et al. 2012, Lundberg, Lebeis et al. 2012, Fitzpatrick, Copeland et al. 2018). These filamentous Gram-positive bacteria are well known for their complex life cycle and their ability to produce a wide range of specialized metabolites, many of which are antibiotics (Donald, Pipite et al. 2022). Increasing evidence suggests that these metabolites also function as cross-kingdom signaling molecules within microbial communities (Krespach, Stroe et al. 2023). The abundance of *Streptomyces* in soils often correlates with resistance to abiotic stresses such as drought and with the suppression of soil-borne fungal pathogens (Cha, Han et al. 2016, Kim, Cho et al. 2019). Recent studies have further elucidated how these bacteria alleviate plant stress through specialized metabolite production (Cha, Han et al. 2016, Yang, Qiao et al. 2023). In addition, *Streptomyces* spp. can modulate plant immunity via mechanisms involving hormonal signaling and iron homeostasis (Conn, Walker et al. 2008, Xiong, Zeng et al. 2024). For instance, suppression of salicylic acid-mediated immunity and iron uptake during drought has been proposed as a key mechanism facilitating *Streptomyces* colonization of *Arabidopsis* roots (Fitzpatrick, Smith et al. 2026). Furthermore, *Streptomyces* can adjust their specialized metabolism in response to changes in the plant metabolome, such as coumarin secretion under stress conditions (Diab, Du et al. 2026).

Despite the recognized importance of *Streptomyces*–plant interactions in stress mitigation, the cross-kingdom signaling processes underlying bacterial colonization and their broader effects on plant physiology and microbiota assembly remain poorly understood (Worsley, Macey et al. 2021, Graindorge, Villette et al. 2022). We previously characterized the rhizospheric strain *Streptomyces* sp. AgN23 and showed that it efficiently colonizes the *Arabidopsis* phyllosphere while activating salicylate and jasmonate-dependent defense pathways required for protection against *Alternaria brassicicola* (Vergnes, Gayrard et al. 2019). Phylogenomic analyses placed AgN23 within the *Streptomyces violaceusniger* genomospecies, a widespread group of soil-dwelling strains commonly associated with plant rhizospheres and characterized by a conserved repertoire of specialized metabolites linked to rhizosphere-associated lifestyles (Gayrard, Nicolle et al. 2023). We further identified galbonolides, a family of polyketides produced by AgN23, as key elicitors of *Arabidopsis* defense through disruption of sphingolipid homeostasis (Nicolle, Gayrard et al. 2024). Using a galbonolide-deficient mutant, we demonstrated that galbonolides reprogram *Arabidopsis* root metabolism by stimulating the production of the indolic alkaloid camalexin, which is essential for AgN23 recruitment in the rhizosphere. However, the broader consequences of this response on rhizosphere microbiota composition remain unclear.

In this study, we investigated two connected aspects of the AgN23–*Arabidopsis* interaction: the plant signalling pathways activated in response to AgN23, and the extent to which these host responses contribute to the structuring of the rhizosphere microbiota. We show that ethylene and salicylic acid signalling contribute to key AgN23-induced responses, including root growth inhibition and the production of antifungal exudates. Metabolomic analyses revealed that AgN23 broadly activates phenylalanine- and tryptophan-derived secondary metabolic pathways in *Arabidopsis* roots, and that this response is abolished in the *npr1* mutant. Furthermore, AgN23 colonization of the rhizosphere was reduced in *npr1* and *ein2* mutants. This indicates that ethylene and salicylic acid signalling act together to support bacterial recruitment. Finally, we demonstrate that AgN23 subtly reprograms both bacterial and fungal rhizosphere communities in an NPR1-dependent manner.

## RESULTS

### NPR1 and EIN2 control AgN23-induced root growth inhibition and antifungal functions

Building on our previous finding that galbonolides mediate the broad metabolomic reprogramming induced by AgN23 in *Arabidopsis* roots (Nicolle, Gayrard et al. 2024), we hypothesized that plant hormone signalling pathways associated with biotic stress responses may coordinate these processes downstream of galbonolide-triggered disruption of sphingolipid biosynthesis. To identify these signalling components, we first used an in vitro inoculation assay in which AgN23 induces root growth inhibition (RGI), a phenotype associated with activation of plant immune responses (Finkel, Salas-González et al. 2020). Under mock conditions, primary root length was comparable between Col-0 and the tested mutants (Figure 1A, 1B). Following AgN23 inoculation, Col-0 and *jar1* showed marked reductions in primary root growth (−23% and −26%, respectively), whereas *ein2* (−1.6%) and *npr1* (−11%) were largely insensitive to the bacterium. These results indicate that salicylic acid- and ethylene-related signalling pathways are required for full expression of the AgN23-induced RGI phenotype.

**Figure 1.**
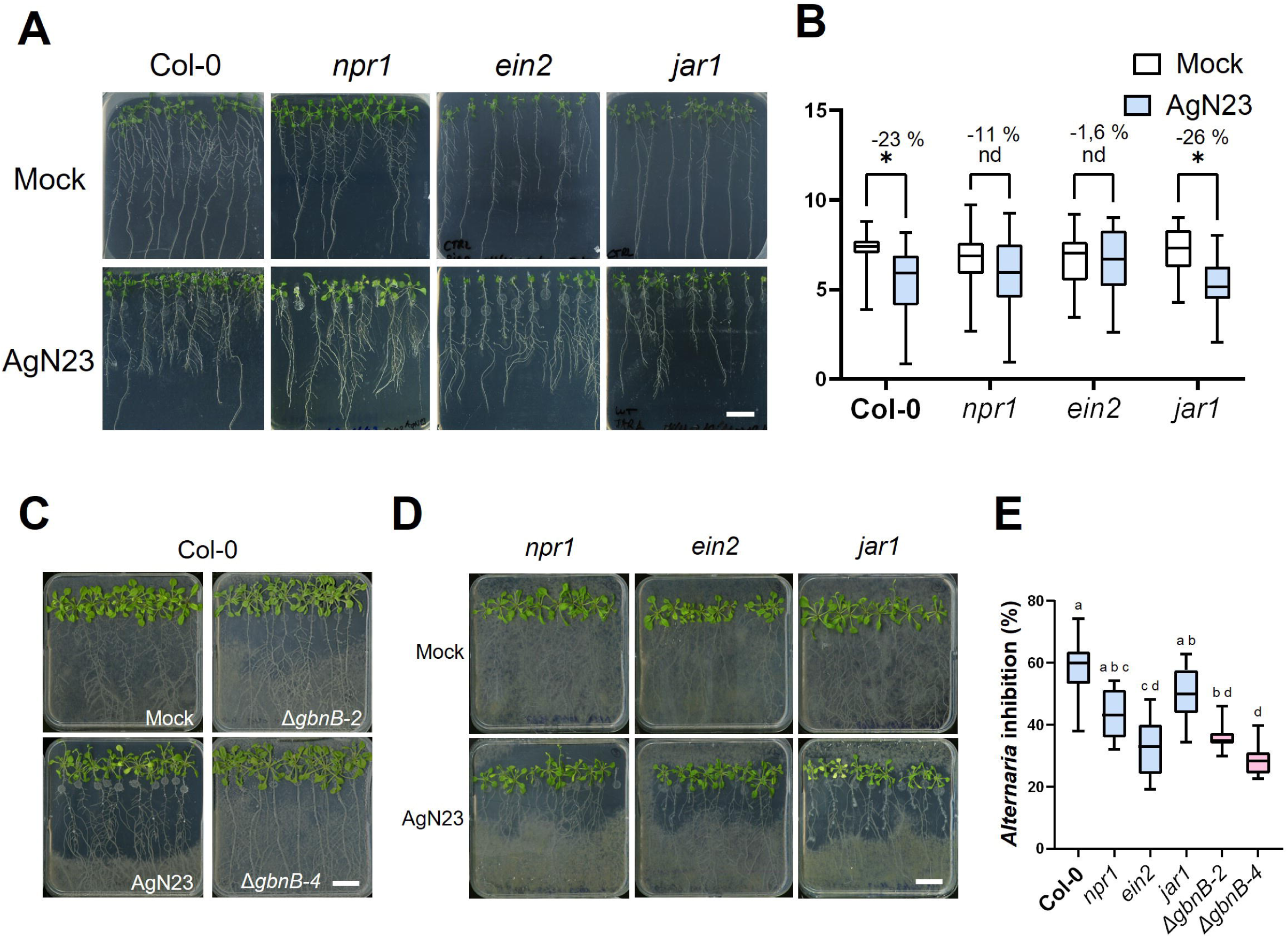
*Arabidopsis* root responses to *Streptomyces* sp. AgN23 and the rhizospheric antifungal activity involve galbonolide production and NPR1- and EIN2-mediated signalling. **(A)** Representative images of *Arabidopsis* thaliana Col-0 and hormonal signalling mutants 10 days after root tip inoculation with *Streptomyces* sp. AgN23 spores (scale bar = 2 cm). **(B)** Primary root length of mock- and AgN23-treated plants at 10 days post-inoculation. Boxplots summarize data from 6–10 independent experiments, each including at least 10 plants per treatment (n > 60). Statistical significance was assessed using a Mann–Whitney test (*P* < 0.05; ns, not significant). Root growth inhibition (%) corresponds to the mean reduction in primary root length between AgN23-treated and mock-treated plants for each genotype. (C) Representative images of *Arabidopsis* seedlings following 10 days of interaction with AgN23 or the galbonolide-deficient mutant (Δ*gbnB*), followed by 4 days of inoculation with *A. brassicicola* (scale bar = 2 cm). **(D)** Representative images of *Arabidopsis* hormonal signalling mutants following 10 days of interaction with AgN23 followed by 4 days of infection with *A. brassicicola*, the upper panel displays mock-treated plants, while the lower panel displays AgN23-inoculated plants (scale bar = 2 cm). **(E)** Quantification of suppression of *A. brassicicola* development on agar medium containing Col-0 and immune signalling mutants inoculated with AgN23 or Δ*gbnB*. Boxplots represent nine independent experiments. Different letters indicate statistically significant differences (ANOVA, *P* < 0.05).

To further characterize the *Arabidopsis*-AgN23 interaction, we next assessed whether AgN23 induces plant-derived antifungal activity around roots. Using an antibiogram-style assay, *Alternaria brassicicola* conidia were sprayed onto plates containing AgN23-inoculated or control plants, and fungal development was monitored over time. In mock-treated Col-0 plants, *A. brassicicola* colonized the entire plate surface, whereas AgN23 treatment reduced fungal spread by approximately 60%. This antifungal effect was weaker with the galbonolide-deficient AgN23 *ΔgbnB* mutant, which reduced fungal growth by less than 40% (Figure 1C, 1E). When the same assay was performed on *npr1*, *ein2*, and *jar1* mutants, AgN23-mediated inhibition of *A. brassicicola* was strongly reduced in *npr1* and *ein2*, but largely retained in *jar1* (Figure 1D, 1E). These findings indicate that fungal suppression around roots depends mainly on galbonolide-induced plant metabolites controlled by NPR1 and EIN2 signalling pathways, rather than on direct microbial antagonism by AgN23.

### NPR1 mediates root metabolomics responses to AgN23

To further characterize the involvement of NPR1 in the root response to AgN23, we performed a comparative global metabolomics analysis of Col-0 and *npr1* roots inoculated with the bacterium. Principal Component Analysis (PCA) of the 2,047 detected variables showed a clear separation between the metabolomes of AgN23-colonized and mock-treated Col-0 roots (Figure 2A). In contrast, the *npr1* metabolome was nearly identical under control and inoculated conditions, and both profiles clustered with untreated Col-0 roots (Figure 2A). These data indicate that the metabolic reprogramming triggered by AgN23 is largely abolished in the *npr1* background.

**Figure 2.**
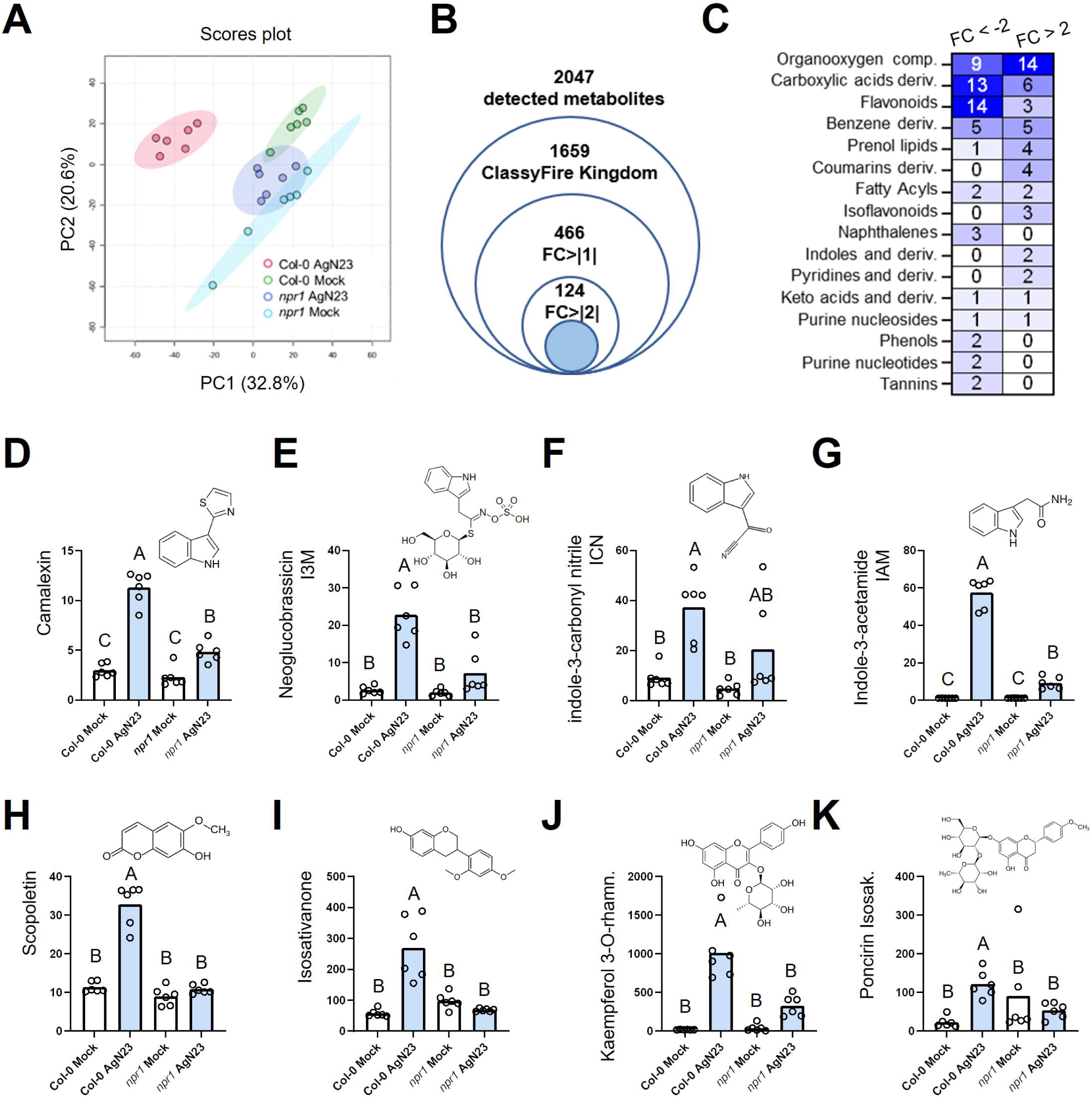
*Streptomyces* sp. AgN23 triggers *Arabidopsis* production of tryptophan- and phenylpropanoid-derived secondary metabolites in a NPR1-dependent manner. **(A)** Principal Component Analysis (PCA) score plot of UHPLC–MS data (n = 2047 variables) from root extracts of *Arabidopsis thaliana* Col-0 and *npr1* mutants under mock conditions and 10 days after inoculation with AgN23. **(B)** Summary of detected variables in the Col-0 root metabolome, including the total number of features detected, those with putative metabolite annotations within the ClassyFire Kingdom ‘Organic Compounds’, and those showing *P* < 0.05 and a |log_2_ FC| > 1 or > 2 in response to AgN23 inoculation. **(C)** Distribution of annotated variables displaying |log_2_ FC| > 2 following AgN23 inoculation across natural product classes defined by ClassyFire. Only classes represented by at least two annotated features showing FC > 2 in Col-0 are displayed. **(D-K)** Hallmark defence-related metabolites produced by *Arabidopsis* roots in an NPR1-dependent manner upon AgN23 inoculation. Metabolites were selected based on their induction in Col-0 following AgN23 inoculation, with fold change (FC) > 2 and *P* < 0.05 in an ordinary one-way ANOVA with Tukey’s multiple comparisons test to assign letters A-C (n = 6 per condition). The plots display the relative ion abundance from LC-MS data.

We next characterized the Col-0 metabolic response by annotating detected variables and assigning them to ClassyFire ontologies. Of the 2,047 detected features, 1,659 received an annotation; among these, 466 showed a log_2_ fold change |log_2_FC| > 1, and 124 displayed stronger regulation (|log_2_FC| > 2) in response to AgN23 (Figure 2B, Supplementary Table S1). Among the features most strongly regulated by AgN23, ClassyFire classification revealed that organooxygen compounds, coumarins, and isoflavonoids were mainly enriched among induced metabolites, whereas carboxylic acids and flavonoids were mostly repressed (Figure 2C). Representative metabolites from these regulated classes were induced by AgN23 in a largely NPR1-dependent manner. These included camalexin (Figure 2D; indoles and derivatives; feature ID 50929_pos in Supplementary Table S1) and neoglucobrassicin (Figure 2E; organooxygen compounds; I3M; 50844_neg), which we previously reported to be induced by AgN23 in a galbonolide-dependent manner(Nicolle, Gayrard et al. 2024). AgN23 also triggered NPR1-dependent accumulation of the cyanogenic glucosinolate indole-3-carbonyl nitrile (Figure 2F; ICN; 51109_neg) and the auxin precursor indole-3-acetamide (Figure 2G; IAM; indoles and derivatives; 50813_pos). Similarly, scopoletin (Figure 2H; coumarin derivatives; 5091_pos) and three isoflavonoid-related metabolites—isosativanone (Figure 2I; 50286_neg), glycosylated kaempferol derivatives (Figure 2J; 50751_pos), and poncirin/isosakuranetin (Figure 2K; 51007_neg) were induced by AgN23 in an NPR1-dependent manner. Together, these results show that AgN23 activates salicylic acid signalling to stimulate phenylalanine- and tryptophan-derived antimicrobial metabolism in *Arabidopsis* roots.

### Elicitation of phenylalanine- and tryptophan-derived secondary metabolism by AgN23 attenuates NPR1-dependent root growth inhibition

We found that NPR1 is required for the full spectrum of *Arabidopsis* responses to AgN23, ranging from primary root growth inhibition to extensive reprogramming of the root metabolome, including the production of defence metabolites derived from phenylalanine and tryptophan metabolism. To disentangle the contribution of these secondary metabolites to the root-growth phenotype, we selected mutants impaired in the corresponding pathways and inoculated them with AgN23 spores. We first examined glucosinolate-deficient mutants affected at distinct steps of the biosynthetic pathway (*cyp79b2/cyp79b3*, *cyp71a12/cyp71a13*, *fox1*, and *cyp82c2*) (Figure 3A, 3B). Ten days after inoculation, most mutants displayed stronger root growth inhibition (RGI) than Col-0: *cyp79b2/cyp79b3* (−44%), *fox1* (−41%), and *cyp82c2* (−43%), compared with −23% in Col-0. The *cyp71a12/cyp71a13* mutant, however, already showed reduced basal root growth under mock conditions (−38% relative to mock-treated Col-0), making its RGI upon AgN23 inoculation (−21%) more difficult to interpret (Figure 3B, 3C). Overall, these data indicate that glucosinolate biosynthesis mitigates AgN23-induced RGI in *Arabidopsis*.

**Figure 3.**
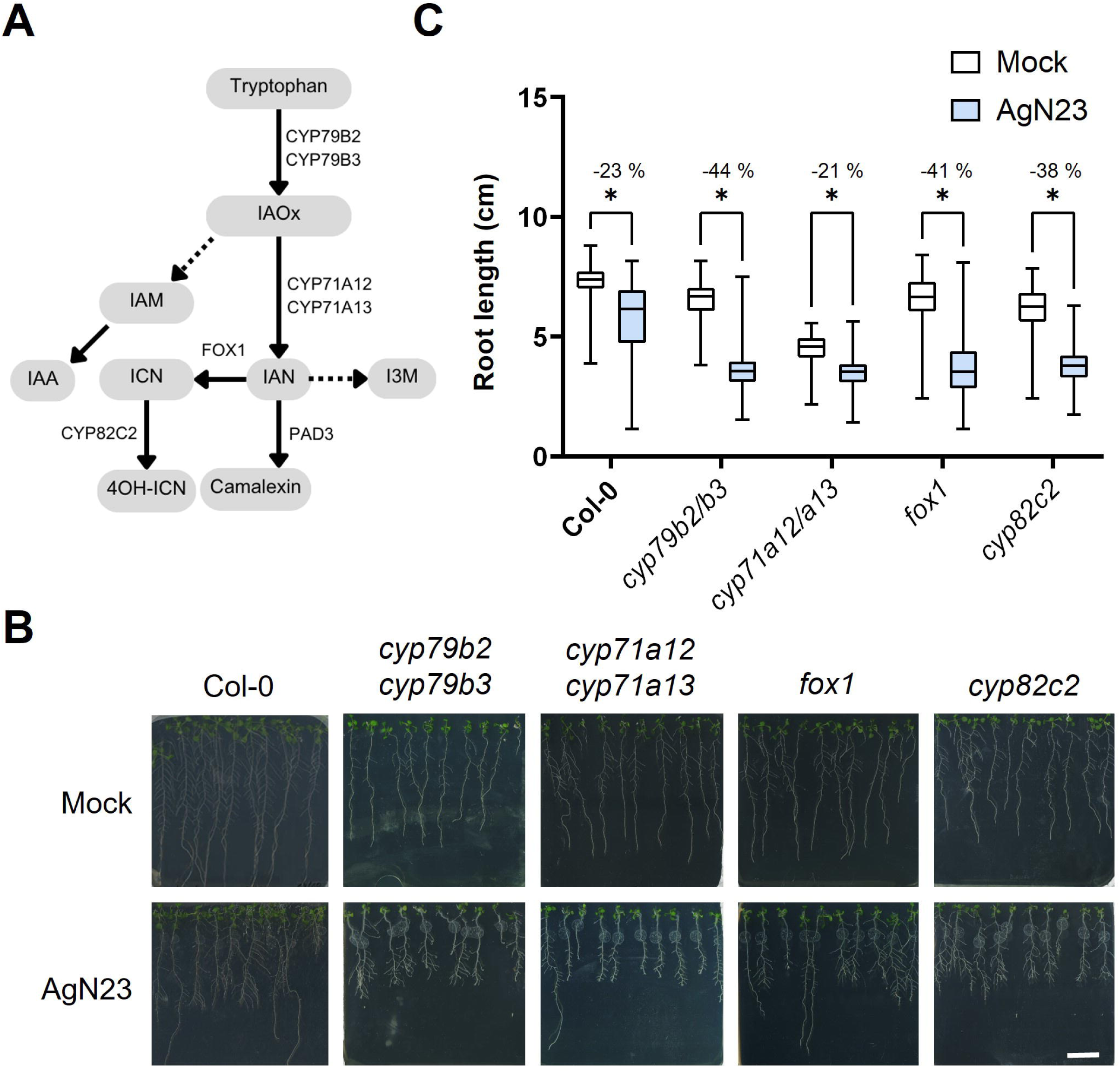
Arabidopsis glucosinolates mitigate AgN23-induced inhibition of root growth. **(A)** Simplified schematic of the biosynthetic pathway of tryptophan-derived metabolites. Grey boxes indicate individual metabolites: (4-OH)-ICN, (4-hydroxy)-indole carbonyl nitrile; IAN, indole-3-acetonitrile; IAOx, indole-3-acetaldoxime; IAM, Indole-3-acetamide. Enzymes catalyzing each step are indicated next to the corresponding arrows. **(B)** Representative images of Arabidopsis thaliana Col-0 and glucosinolate mutants grown under mock conditions or colonized by AgN23. Photographs were taken 10 days after inoculation with spores applied to the root apex (scale bar = 2 cm). **(C)** Primary root length of glucosinolate-deficient mutants. Boxplots summarize data from at least six independent experiments, each including ≥10 seedlings per treatment (n > 60). Statistical differences between mock and AgN23 treatments were assessed within each genotype using a Mann– Whitney test; \**P* < 0.05. Inhibition percentages were calculated by comparing AgN23-treated and mock conditions for each genotype.

We next investigated mutants impaired in flavonoid (*tt4*) and coumarin (*myb72-2*, *f6’h1*) biosynthesis (Figure 4A). Under mock conditions, all mutants produced shorter roots than Col-0, with reductions ranging from −19% to −26% (Figure 4B). Following AgN23 inoculation, these mutants exhibited stronger RGI than Col-0: *tt4* (−43%), *myb72-2* (−40%), and *f6’h1* (−34%). These results suggest that, as observed for glucosinolates, AgN23-induced activation of flavonoid and coumarin pathways helps buffer the RGI caused by *Streptomyces*. This conclusion contrasts with the phenotype of the *npr1* mutant, in which these secondary metabolic pathways are not induced, yet strong RGI is not observed. A plausible explanation is that *npr1* displays a broader collapse of exudate-related metabolism, which may reduce AgN23 growth in the rhizosphere and thereby limit its capacity to trigger RGI.

**Figure 4.**
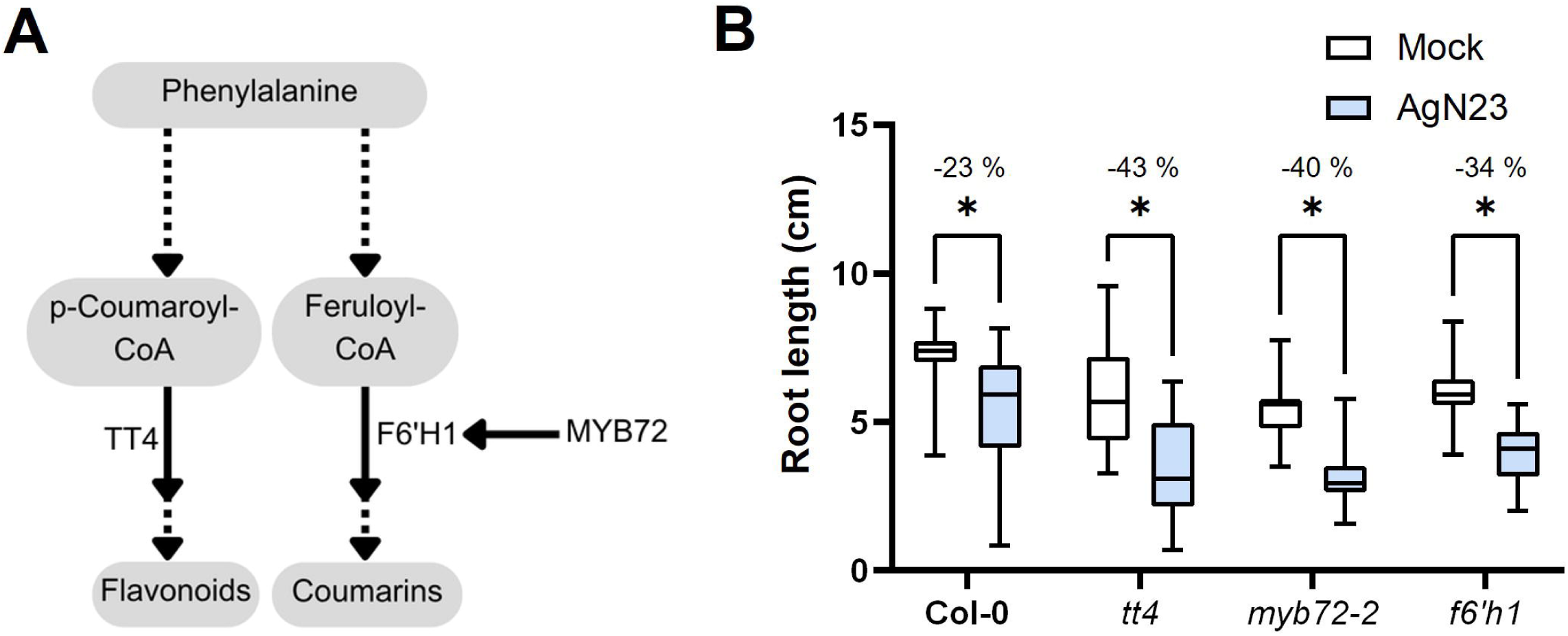
Phenylpropanoids mitigate AgN23-induced root growth inhibition of *Arabidopsis* thaliana. **(A)** Simplified schematic of the flavonoid and coumarin biosynthetic pathways in *Arabidopsis*. Enzymes catalyzing individual metabolic steps are indicated next to the corresponding arrows. **(B)** Primary root length of seedlings colonized or not by AgN23, measured 10 days after inoculation. Boxplots represent data from 6–10 independent experiments, each including at least 10 seedlings per treatment (n > 60). Statistical differences between mock and AgN23 treatments were assessed using a Mann–Whitney test; \**P* < 0.05. Inhibition percentages were calculated by comparing AgN23-treated and mock conditions for each genotype.

### NPR1 and EIN2 contribute to rhizosphere recruitment of AgN23

We previously observed that AgN23 rhizosphere colonization is severely impaired in the camalexin-deficient *pad3* line, suggesting that plant immune activation promotes bacterial development in the rhizosphere (Nicolle, Gayrard et al. 2024). To test whether NPR1 contributes to this process, we performed soil inoculation assays with *npr1* plants and extended the analysis to the *ein2* and *jar1* mutants examined above. Ten plants per genotype were uprooted and pooled in pairs for rhizosphere sampling and DNA extraction, while cleaned roots were processed separately to quantify AgN23 levels in the rhizosphere. Using the 2^-ΔCt^ method, we quantified a single-copy AgN23 marker by qPCR in rhizosphere DNA and normalized its abundance against V3–V4 16S rRNA amplicons representing the total bacterial community. In Col-0, AgN23 reached approximately 30 ppm relative to 16S copies, corresponding to a qPCR signal of 3 × 10^-5^ (Figure 5A) with high repeatability among samples. Across mutant backgrounds, AgN23 relative abundance was significantly reduced in *npr1* and *ein2*, reaching 11 and 8 ppm, respectively, corresponding to an approximately 30% reduction in AgN23 enrichment in both mutant rhizospheres and large variability among the samples. These data suggest that salicylic acid and ethylene signalling are required for optimal recruitment of *Streptomyces* AgN23 and for the establishment of typical plant responses to colonization, including root growth inhibition and antifungal exudate production documented above.

**Figure 5.**
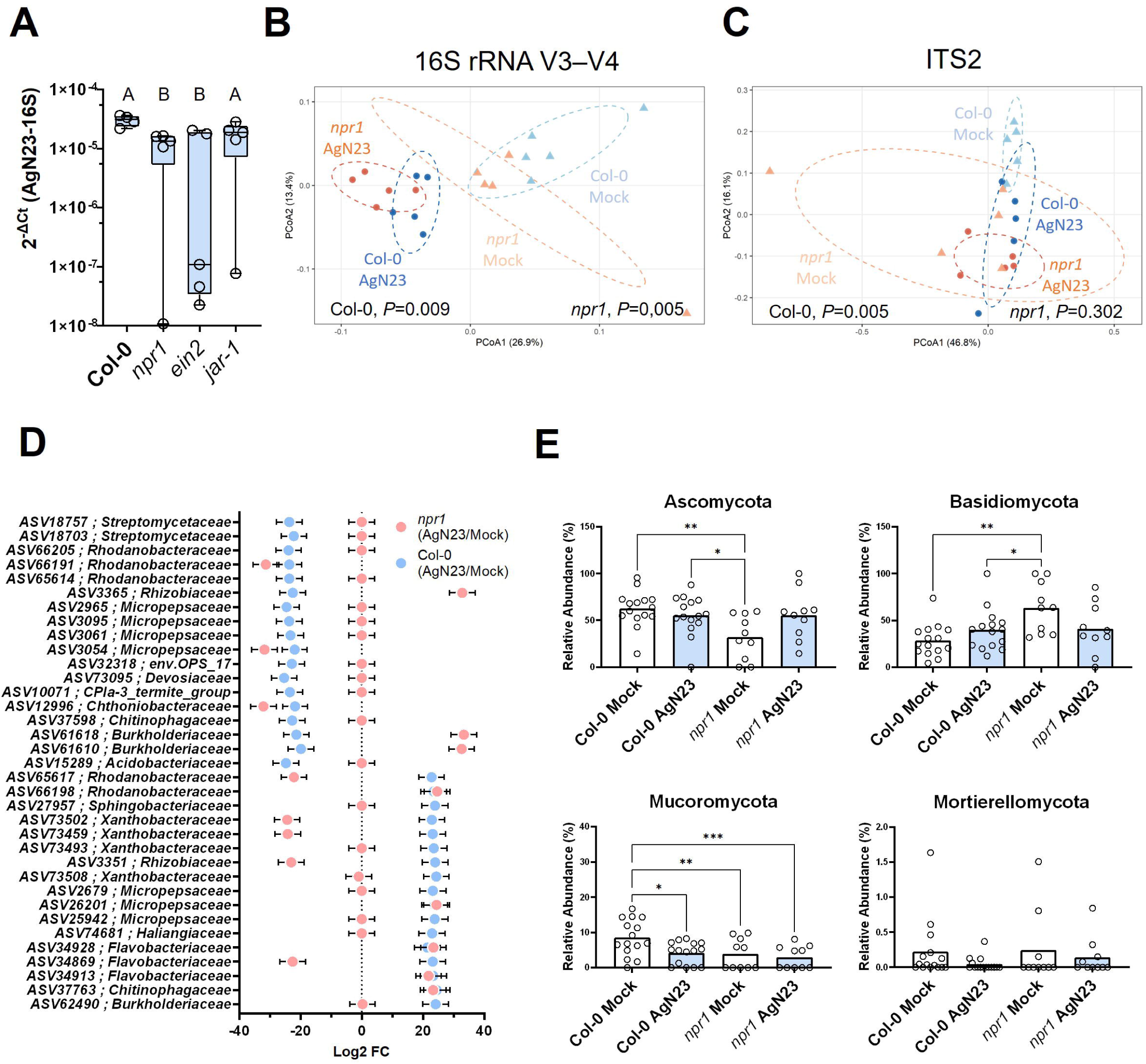
Colonization of the rhizosphere and reprogramming of the microbiota by AgN23 require NPR1. (A) Quantification of AgN23 genomic DNA relative to 16S rRNA by qPCR in the rhizosphere 6 weeks after soil inoculation. Boxplots represent data from 10 plants per condition; whiskers indicate minimum and maximum values, and the central line shows the median. Different letters (A–B) denote statistically distinct groups based on Kruskal–Wallis multiple comparisons. (B) Principal component analysis (PCA) of bacterial community beta diversity based on Bray–Curtis distances using the 16S rRNA V3–V4 region. PERMANOVA was performed to compare mock and AgN23-treated rhizospheres in both Col-0 and npr1 backgrounds; corresponding P-values are indicated (n = 5). (C) Principal component analysis (PCoA) of fungal community beta diversity based on Bray–Curtis distances using the ITS2 marker. PERMANOVA was conducted to compare mock and AgN23-treated rhizospheres in both Col-0 and npr1 backgrounds; P-values are indicated (n = 5). (D) Bacterial taxonomic assignment at the family level of amplicon sequence variants (ASVs) showing significant differential abundance) in AgN23-treated rhizosphere of Col-0 plants (log_2_ fold change, *P* < 0.001. (E) Relative abundance (%) of the four most abundant fungal phyla following AgN23 inoculation in Col-0 and npr1. Statistical significance was assessed using Tukey’s multiple comparisons test (\**P* < 0.05, \*\**P* < 0.01, \*\*\**P* < 0.001).

### NPR1 largely drives AgN23-mediated modulation of the rhizosphere microbiota

The immunity-stimulating and antimicrobial activities of AgN23 documented previously and in this study led us to hypothesize that AgN23 colonization may modulate the composition of the *Arabidopsis* rhizosphere microbiota. To separate direct effects of AgN23 on surrounding microorganisms from indirect effects mediated by host responses, we used metabarcoding to compare bacterial and fungal communities in Col-0 and in the *npr1* mutant, which we showed above to be largely compromised in its responses to the bacterium. Bacterial communities were analysed using the 16S rRNA V3–V4 region, whereas fungal communities were profiled using ITS2, in rhizosphere samples from Col-0 and *npr1* plants grown under mock or AgN23-inoculated conditions.

Using Bray–Curtis dissimilarity to assess microbial community composition, we found that AgN23 inoculation significantly altered the bacterial communities of both Col-0 and *npr1* rhizospheres (Figure 5B; PERMANOVA, P = 0.003 and P = 0.009, respectively). However, bacterial communities also differed significantly between mock-treated Col-0 and *npr1* plants (P = 0.045), indicating that *npr1* plants assemble a dysbiotic rhizosphere bacteriota even in the absence of AgN23. Consistently, AgN23-treated communities from the two genotypes remained significantly distinct (P = 0.011), showing that the *npr1* response to AgN23 does not recapitulate the microbiota reprogramming observed in wild-type Col-0 plants.

We next used DESeq2 to identify AgN23-responsive bacterial ASVs (Figure 5D). Among 7,189 detected ASVs, observed richness ranged from 300 to 450 ASVs across mock and AgN23-treated Col-0 and *npr1* samples, with no significant differences between groups. DESeq2 analysis identified a total of 54 ASVs significantly regulated by AgN23 (*P* < 0.001) either in Col-0 (35 ASVs) or in *npr1* with 35 ASVs as well. Among the 35 AgN23-responsive ASVs identified in each genotype, 16 overlapped between genotypes. However, only eight of these 16 ASVs were regulated in the same direction, whereas the remaining eight displayed opposite responses in Col-0 and *npr1* (Supplementary Table S3). Shared AgN23-induced ASVs included two *Flavobacteriaceae* (ASV34913, ASV34928), one *Micropepsaceae* (ASV26201), one *Rhodanobacteraceae* (ASV66198), and one *Chitinophagaceae* (ASV37763), whereas shared repressed ASVs belonged to *Chthoniobacteraceae* (ASV12996), *Micropepsaceae* (ASV3054), and *Rhodanobacteraceae* (ASV66191). These shared responses define an NPR1-independent component of AgN23-mediated microbiota reprogramming.

In contrast, several AgN23-responsive ASVs were regulated only in Col-0 and thus in an NPR1-dependent manner. Twelve ASVs induced by AgN23 in Col-0 belonged notably to *Flavobacteriaceae* (ASV34869), *Micropepsaceae* (ASV25942, ASV2679), *Rhodanobacteraceae* (ASV65617), and *Xanthobacteraceae* (ASV73508, ASV73493, ASV73459, ASV73502). Many of these ASVs were less abundant in AgN23-treated *npr1* rhizospheres than in Col-0, suggesting that NPR1 is required for their recruitment by AgN23 (Supplementary Table S3). Conversely, among the 15 ASVs depleted by AgN23 in Col-0, we detected *Burkholderiaceae* (ASV61610, ASV61618), *Rhodanobacteraceae* (ASV65614, ASV66205), *Micropepsaceae* (ASV3054, ASV3061, ASV3095, ASV2965) and *Streptomycetaceae* (ASV18703, ASV18757), most of these ASVs were constitutively depleted in mock-treated *npr1* plants (Figure 5D, Supplementary Table S3). Overall, AgN23-driven reprogramming of the rhizosphere bacteriota is therefore largely, but not completely, dependent on functional NPR1.

Notably, AgN23 also regulated ASVs in the *npr1* background that were not regulated in Col-0, including eight induced and eleven repressed ASVs belonging mainly to *Burkholderiaceae*, *Flavobacteriaceae*, *Micropepsaceae*, and *Rhodanobacteraceae*. This indicates that AgN23 can also modulate rhizosphere communities independently of the typical NPR1-dependent host metabolic response, potentially through direct microbial competition, including competition with a *Streptomycetaceae* ASV that is initially overrepresented in the *npr1* mock rhizosphere (ASV18697). Taken together, these results suggest that AgN23 promotes its own rhizosphere establishment partly at the expense of other *Streptomyces*-related taxa, while concomitantly favouring the recruitment of plant growth-promoting rhizobacteria such as *Flavobacteriaceae*.

We used the same rhizosphere samples to assess the impact of AgN23 on fungal communities. Bray–Curtis dissimilarity analysis showed that AgN23 significantly altered fungal community composition in Col-0 (Figure 5C; PERMANOVA, P = 0.003), but not in *npr1* (P = 0.261). A total of 379 fungal ASVs were detected, with observed richness ranging from 50 to 75 ASVs across mock and AgN23-treated Col-0 and *npr1* samples. However, DESeq2 analysis did not identify any fungal ASV showing significant AgN23-dependent changes in either genotype (*P* < 0.001).

We therefore examined AgN23-driven changes in fungal community composition at the phylum level (Figure 5E). In Col-0 plants, AgN23 inoculation did not significantly affect the relative abundance of *Ascomycota* or *Basidiomycota*, the two dominant fungal phyla in the rhizosphere. In contrast, *Mucoromycota* was strongly depleted following AgN23 treatment. This depletion was not observed in the *npr1* background, consistent with the PERMANOVA results. These findings indicate that AgN23-mediated restructuring of the rhizosphere mycobiota depends on intact SA/NPR1 signalling in *Arabidopsis*, rather than on direct antifungal activity of the bacterium, such as galbonolide-mediated inhibition.

## DISCUSSION

*Streptomyces* spp. are well-established contributors to plant stress resilience, largely through the production of specialized metabolites that support root development, microbial colonization, and stress adaptation. For example, pteridic acid promotes rhizogenesis under drought conditions, illustrating the close link between specialized metabolism, root-associated lifestyles, and plant-beneficial effects. AgN23 provides a particularly relevant example of this paradigm. This strain produces galbonolides, a family of polyketides that target plant inositol phosphoceramide synthase, disrupt sphingolipid homeostasis, and also display antifungal activity (Nicolle, Gayrard et al. 2024). We previously demonstrated that galbonolides produced by *Streptomyces* sp. AgN23 elicit plant immunity, thereby promoting AgN23 colonization of the rhizosphere and ultimately enhancing resistance to biotic stress.

Here, we investigated the downstream events of AgN23 rhizosphere colonization on the host plant and its microbiota. Using the *npr1* mutant, we demonstrate that SA signaling is required downstream of AgN23-derived galbonolides to mediate root growth inhibition and to induce antifungal responses associated with tryptophan- and phenylalanine-derived metabolism. In addition, the induction of key root exudates is compromised in *npr1*, highlighting NPR1 as a central regulator of root metabolic profile. Consistent with established models, the similar phenotypes observed in *ein2* and *npr1* emphasize that ethylene and salicylate signaling triggered by AgN23 cooperatively regulate root immunity and development (Růžička, Ljung et al. 2007). By contrast, JAR1-dependent jasmonate signaling is known to antagonize NPR1 responses, consistent with the wild-type behavior of this mutant upon AgN23 inoculation (Ahn, Lee et al. 2007, Leon-Reyes, Spoel et al. 2009). Altogether, these results identify NPR1-dependent salicylic acid signaling as a central upstream component of the plant signaling cascade activated by galbonolides.

AgN23 triggers extensive NPR1-dependent metabolic rewiring, including activation of phenylalanine- and tryptophan-derived pathways, as well as primary metabolites belonging to “Organooxygen compounds” and “Carboxylic acid derivatives” ClassyFire categories which encompass simple sugars and amino acids (Tsai, Tang et al. 2026, Yu, Liu et al. 2019, O’Banion, Jones et al. 2023, Bhattacharyya, Pablo Clint et al. 2025). These changes likely establish a metabolic environment favorable to AgN23 colonization. Paradoxically, the loss of glucosinolate, flavonoid, or coumarin biosynthesis enhances root growth inhibition in the corresponding mutants, whereas this response is abolished in *npr1*, which is impaired across all of these pathways. In addition, we previously reported that *pad3* displays wild-type root growth in the presence of AgN23. However, unlike camalexin, which is inducible, many tryptophan- and phenylalanine-derived metabolites are constitutively present, and disruption of their biosynthesis can alter root development. CYP79B2 and CYP79B3 convert tryptophan into indole-3-acetaldoxime (IAOx), the precursor of both auxin (IAA) and indole glucosinolates (Hull, Vij et al. 2000). Consistently, CYP79B2 overexpression leads to auxin overproduction phenotypes, whereas *cyp79b2/cyp79b3* mutants display reduced IAA levels and growth defects reflecting auxin deficiency (Zhao, Hull et al. 2002).

The detection of indole-3-acetamide (IAM) in our metabolomic dataset further supports crosstalk between auxin and stress pathways, as IAM directly inhibits root growth and contributes to both auxin biosynthesis and abscisic acid-mediated stress responses(Malka and Cheng 2017, Pérez-Alonso, Ortiz-García et al. 2021, Sánchez-Parra, Pérez-Alonso et al. 2021). By contrast, components of the indole and glucosinolate pathways that act further downstream, such as CYP71A12 and PAD3, primarily contribute to the production of defense metabolites rather than to root development changes (Koprivova, Schuck et al. 2019, Pastorczyk, Kosaka et al. 2020, Koprivova, Schwier et al. 2023). Notably, similar regulatory roles are observed for flavonoids and coumarins, which influence root architecture and auxin transport (Lupini, Araniti et al. 2014, Chapman and Muday 2021).

AgN23-induced metabolic reprogramming has clear consequences for rhizosphere microbiota structure. Our data reveal a decrease in *Streptomycetaceae* and an enrichment of *Flavobacteriaceae* within bacterial communities, alongside a reduction in *Mucoromycota* within fungal communities, while *Ascomycota* and *Basidiomycota* remain largely unaffected. The reduction in *Streptomycetaceae* suggests competitive exclusion among closely related taxa, potentially driven by niche occupation or specialized metabolite production (Zhao, Bertolli et al. 2024). In contrast, AgN23 appears to recruit *Flavobacteriaceae* adapted to *Arabidopsis* root defense metabolism and potentially involved in fungal pathogen suppression through antimicrobial specialized metabolites (Carrión, Perez-Jaramillo et al. 2019, Lidbury, Borsetto et al. 2020, Prout, Williams et al. 2024, Zhao, Bertolli et al. 2024). *Flavobacteriaceae* are also hallmark phosphate-solubilizing bacteria; therefore they may also contribute to host phosphorus nutrition (Lidbury, Borsetto et al. 2020). Such phosphate provisioning by the bacteriota promoted by AgN23 is consistent with the selective depletion of *Mucoromycota* from the *Arabidopsis* mycobiota. Indeed, these fungi are thought to establish endophytic exchange structures that supply nutrients such as phosphate to the host and may be reduced when plants have access to alternative nutrient acquisition pathways (Prout, Williams et al. 2024).

The NPR1-dependent metabolic reprogramming triggered by AgN23 has direct consequences for its recruitment in the rhizosphere, and roots colonized by AgN23 harbor altered bacterial and fungal communities. Thus, NPR1 not only controls plant defense responses but also shapes microbiota assembly. Although AgN23 possesses the capacity to produce antimicrobial compounds such as galbonolides, our data indicate that microbiota shifts are primarily driven by the plant defense–activating properties of these polyketides rather than by direct antifungal effects. These data are consistent with in vitro inoculation assays performed on *Arabidopsis* using *Alternaria brassicicola*. Altogether, these findings indicate that NPR1 integrates root development, immune signaling, and microbial recruitment, acting as a key regulator of plant microbiota homeostasis and the extended plant immune system. Accordingly, AgN23 primarily acts through host reprogramming rather than direct antimicrobial effects.

In conclusion, we demonstrate that AgN23 reprograms host physiology by targeting NPR1-dependent metabolic pathways. This host-mediated mechanism enhances AgN23 rhizospheric fitness while simultaneously reshaping root architecture and microbiota function. Although this cascade results in mild growth inhibition under our experimental conditions, the strong activation of phenylalanine- and tryptophan-derived secondary metabolism is likely to enhance plant resilience to diverse environmental stresses, including nutrient limitation, fungal pathogens, and insect herbivory (Liu, Jiang et al. 2010, Harbort, Hashimoto et al. 2020, Alfonso, Stahl et al. 2021, Nguyen, Trotel-Aziz et al. 2022).

## MATERIALS AND METHODS

### Plant material, growth conditions and phenotyping

Seeds of *Arabidopsis thaliana* accession Col-0 (N1092) were obtained from the Nottingham Arabidopsis stock center (NASC) as well as some of the mutants listed in Table 1. Other mutants were provided by collaborators as detailed in Table 1. For soil inoculation assays with *Streptomyces* sp. AgN23, the *Arabidopsis* seeds were sown in potting soil (PROVEEN; Bas Van Buuren B.V., Holland) and cultivated in a growth chamber under 16-hour photoperiod and 22°C. The spore inoculation assay in soil was conducted as described previously (Nicolle, Gayrard et al. 2024). Briefly, 58 g of potting soil inoculated with AgN23 spore inoculum at 10^5^ CFU/g was distributed in pots. Similarly, *in vitro* inoculation of AgN23 on *Arabidopsis* was conducted as described previously (Nicolle, Gayrard et al. 2024). Briefly, seeds were surface sterilised and placed for 2 days in phytotron chamber on solid Murashige and Skoog (MS, Sigma) at 4.4 g/L amended with 1% sucrose. The seeds were allowed to sprout before being transferred to sucrose-free MS medium and then inoculated with AgN23 spores. The image acquisition was performed with an Epson Expression 11000 XL scanner. Primary root lengths and rosette area were measured with ImageJ software (v. 1.51k).

**Table 1:**
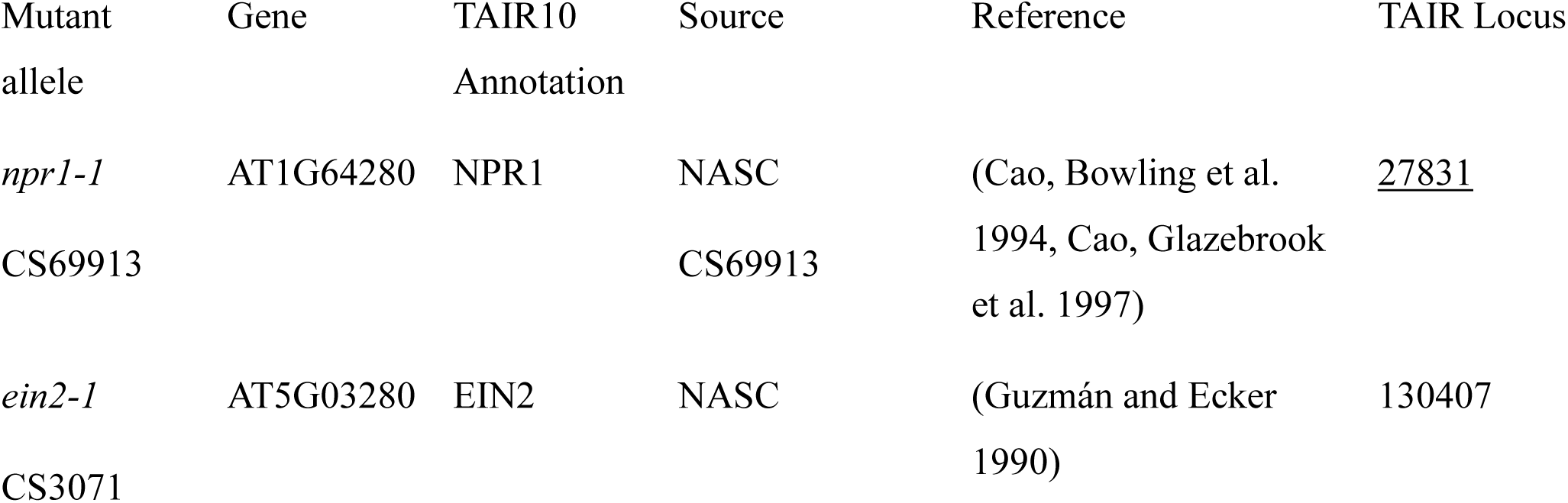
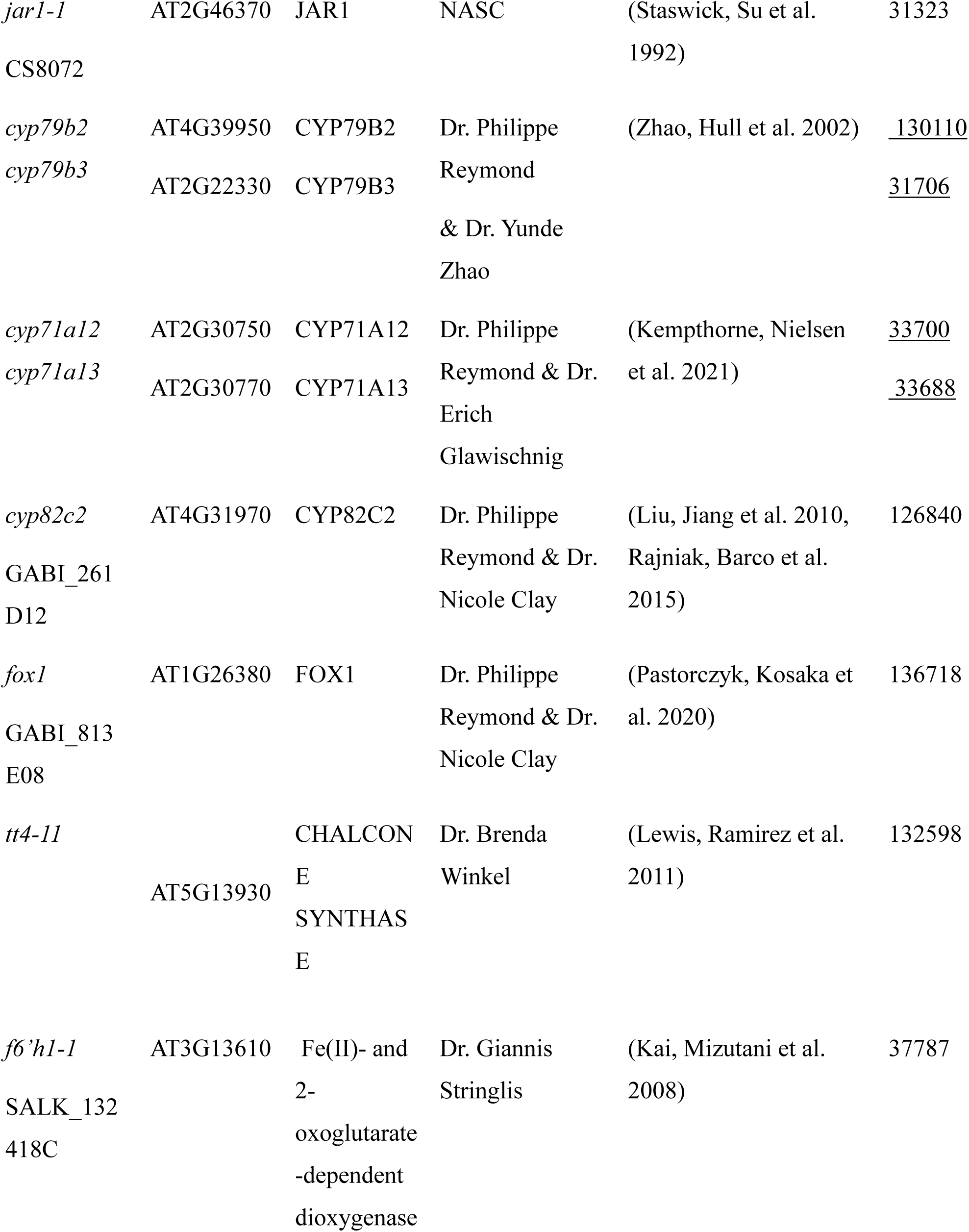
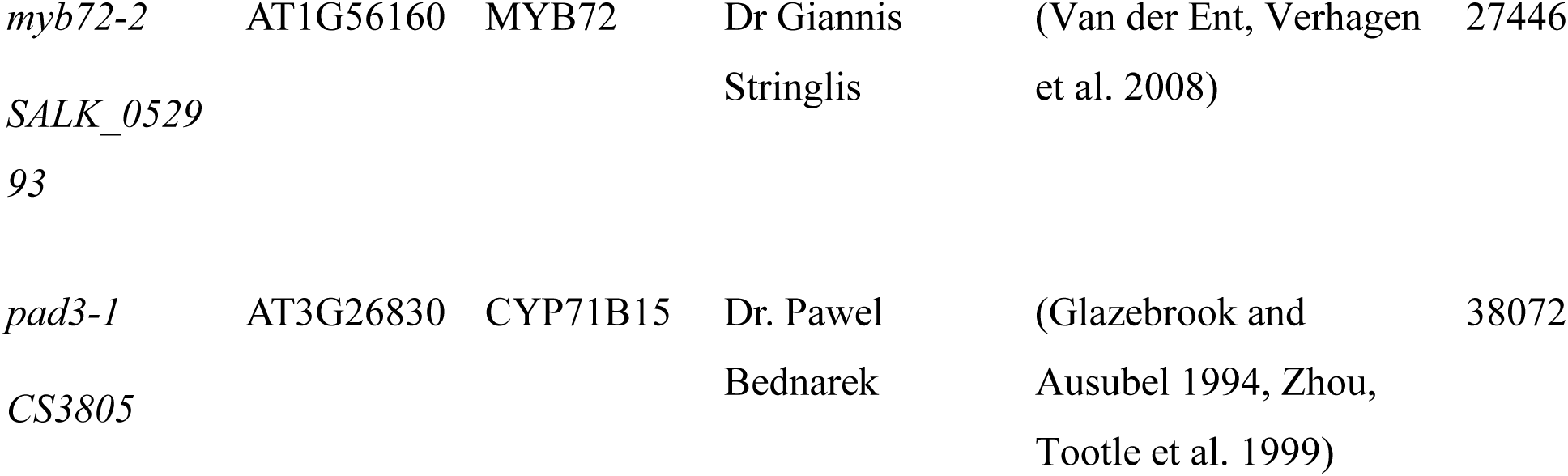

### *Streptomyces* sp. AgN23 cultivation and inoculation

The culture of *Streptomyces* sp. AgN23 and the galbonolides biosynthesis-deficient Δ*gbnB* mutants have been conducted as described previously (Nicolle, Gayrard et al. 2024). Briefly, spore inoculum was obtained from Soy Flour Mannitol plates incubated for two weeks at 28°C in darkness before filling them with 10 mL of sterile ultrapure water. The mycelium was thoroughly scraped with a sterile spreader and the resulting solution was vortexed, then filtered in a 20 mL syringe filled with sterile cotton wool. The resulting suspension was centrifuged at 4,200 rpm for 10 minutes. The supernatant was discarded, the pellet re-suspended in sterile ultrapure water, and the resulting suspension was adjusted to 10^5^ CFU/mL. This spore inoculum was used for subsequent inoculation of *Arabidopsis* in pot and *in vitro*.

### *Alternaria brassicicola* inoculation

*Alternaria brassicicola* strain Abra43 (Sellam, Dongo et al. 2007) was maintained on potato dextrose agar as previously described (Vergnes, Gayrard et al. 2020). *A. brassicicola* spores were prepared as a suspension at 2 × 10⁵ spores mL⁻¹. A volume of 500 µL of the suspension was sprayed per Petri dish, corresponding to 1 × 10⁵ spores per plate. Inoculation was performed using 10 uniform sprays (“sprays”) from a 10 mL glass atomizer to ensure complete coverage of the agar surface. The same sprayer was used for all plates within an experiment to maximize treatment homogeneity. Plates were scanned 4 days after *A. brassicicola* inoculation. For each plate, the fungal growth area was manually delineated in ImageJ using the ROI tool and quantified from pixel counts. The percentage of AgN23-induced inhibition was then calculated by comparing the *A. brassicicola*-covered area with the total plate surface.

### DNA extraction from roots and rhizosphere samples

Root samples (pool of two plants per sample) were carefully washed to remove loosely attached particles, finely chopped, and thoroughly homogenized by a grinding step in 2 mL microcentrifuge tubes containing one steel bead (3 mm) and two glass beads (2 mm). Mechanical disruption was performed using a Retsch MM400 bead mill at 30 Hz, for five cycles of 1 min. The first two grinding cycles were performed at room temperature with thawed samples, followed by three additional cycles with samples frozen in liquid nitrogen to enhance tissue disruption. DNA was extracted using the ZymoBIOMICS™ 96 DNA Kit (Zymo Research), following the manufacturer’s instructions. In brief, the resulting ground roots were resuspended in 400 µL of lysis buffer and transferred to the 96-well plates supplied with the ZymoBIOMICS kit. An additional 250 µL of lysis buffer was added to reach the final volume recommended by the manufacturer (650 µL). Rhizosphere pellets were resuspended in 2 mL of 1× PBS and homogenized. An aliquot of 100 µL of the suspension was transferred to the ZymoBIOMICS 96-well plate tubes, followed by the addition of 550 µL of lysis buffer. Samples were then disrupted using a FastPrep-96 instrument (MP Biomedicals) for four cycles of 1 min at maximum speed (1800 CPM). DNA concentrations were determined using a NanoDrop spectrophotometer. All DNA samples were normalized to a final concentration of 2 ng/µL for downstream qPCR and metabarcoding analyses.

### AgN23 quantification and high-throughput sequencing of 16S rRNA and ITS2 amplicons

AgN23 abundance in the rhizosphere was quantified by qPCR using previously designed AgN23-specific primers and normalized to the bacterial 16S rRNA V3–V4 signal using the 2^-ΔCp^ method (Supplementary Table S3)(Nicolle, Gayrard et al. 2024). For fungal and bacterial metabarcoding, the sequencing libraries were prepared following the Illumina two-step PCR protocol and normalized on SequalPrep plates (ThermoFisher™). The bacterial and fungal primer sets, targeting the V3–V4 region of the 16S rRNA gene (16S rDNA) and the ITS2 region respectively, are listed in Supplementary Table S2. Amplicon paired-end sequencing was carried out at the GeT-Biopuces platform (Genotoul, Toulouse, France) on an Illumina MiSeq instrument with v2 chemistry, generating 2 × 250-bp reads in accordance with the manufacturer’s instructions. Raw reads in FASTQ format were analysed with QIIME2 (v. 2021.4). Paired-end reads were merged, quality-filtered, dereplicated, and screened for chimeras using the DADA2 pipeline, which also resolved the representative amplicon sequence variants (ASVs). Taxonomy was assigned to the 16S and ITS2 sequences with a pre-trained naive Bayes classifier, using the SILVA 138 and UNITE v8 reference databases, respectively.

### Preparation of samples for mass spectrometry analysis

The sample preparation for metabolomic analysis of *Arabidopsis* root was performed as described previously with the following modifications (Nicolle, Gayrard et al. 2024). Briefly, 100 mg of *Arabidopsis* flash frozen samples were ground with FastPrep system 2 times *via* bead beating into lysing matrix D 2 mL tubes (MP Biomedicals) containing 1 mL of extraction buffer (CH3OH: C3H8O: H2O: CH₂O₂ = 40: 40: 19.5: 0.5). The supernatants were recovered by centrifugation and concentrated in a Speedvac vacuum concentrator (Thermo Scientific). Samples were then resuspended in 1 mL of injection buffer (50% CH3OH 50% H_2_O) and filtered through 750 µL nonsterile micro-centrifugal filters (PTFE; 0.2 µm; Thermo Scientific) and introduced into HPLC-certified vials. An aliquot of each sample from the same extraction series was pooled together for quality control (QC).

### UHPLC-HRMS, features annotations and statistical analysis of metabolomics datasets

The analytical parameters for sample injection in Ultra High Performance Liquid Chromatography (UHPLC) and High Resolution Mass Spectrometry (HRMS) analysis were the same as described in Nicolle et al. (2024).

The raw data were processed with MS-DIAL version 4.70 for mass signal extraction between 100 and 1,500 Da from 0.5 to 10.6 minutes, respectively (Tsugawa, Cajka et al. 2015, Fraisier-Vannier, Chervin et al. 2020). MS1 and MS2 tolerances were set to 0.01 and 0.025 Da in the centroid mode. The optimized detection threshold was set to 5 × 10^5^ concerning MS1 and 10 for MS2. Peaks were aligned on a QC reference file with a retention time tolerance of 0.15 minutes and a mass tolerance of 0.015 Da. Minimum peak height was set to 70% below the observed total ion chromatogram (TIC) baseline for a blank injection. Peak annotation was performed with MS-FINDER version 3.52 (Tsugawa, Kind et al. 2016). The MS1 and MS2 tolerances were, respectively, set to 10 and 20 ppm. Formula finders were only processed with C, H, O, N, and S atoms. Taxonomically focused databases (DBs) for Arabidopsis, Brassicaceae, and Streptomyces were generated from COCONUT v2 (Chandrasekhar, Rajan et al. 2025). The internal generic DBs from MS-FINDER used were KNApSAcK, PlantCyc, NANPDB, UNPD, COCONUT, and CheBI.

Output files from MS-DIAL and MS-FINDER were imported into MS-Net (Pereira Francisco, Duthen et al. 2026). Among the top 10 *in silico* candidates per feature, those matching taxonomic criteria (genus, family) were elevated to Level 3a. Features were filtered using RT clustering (ΔRT ≤ 0.01 min). Within each cluster, the top 2 features by network degree and the top 2 by peak intensity were retained. The Mass Spectral Similarity network was constrained to edges with cosine similarity ≥ 0.7 and ΔRT ≤ 8 min between connected nodes. High-confidence annotations (Levels 1, 2a, 3a) seeded the MSS network for iterative annotation propagation. For each feature pair, candidate structures were ranked using a weighting parameter set to α = 0.3, prioritizing structural-spectral evidence (70%) over *in silico* ranking (30%). The top 5 candidates by Link Score were retained per feature before looping through the entire MSS network. Redundant annotations with identical InChIKey identifiers and Pearson correlation among samples > 0.7 were consolidated by selecting the candidate with the highest mean peak height. Positive and negative mode feature lists were merged using ΔRT ≤ 0.05 min, Δm/z ≤ 0.002 Da, and a minimum Pearson correlation ≥ 0.6 across sample intensities. Final annotations were enriched with chemical ontology classifications from ClassyFire (kingdom, superclass, class, subclass) and NPClassifier (pathway, superclass, class). Database identifiers (PubChem CID, KEGG, HMDB, ChEBI) were retrieved using the Chemical Translation Service.

## DATA AVAILABILITY

Datasets generated or analysed during this study are included in this published article (and its Supplementary Table files). The LC-HRMS chromatograms of *Arabidopsis* responses to AgN23 CME are available in the Zenodo repository (8421008 and 11621475). The rhizosphere metabarcoding dataset of rhizosphere is available at NCBI under the BioProject PRJNA1481870.

## ACKNOWLEDGEMENTS

We thank Dr. Philippe Reymond, Dr. Yunde Zhao, Dr. Erich Glawishnig, Dr. Nicole Clay, Dr. Brenda Winkel, and Dr. Giannis Stringlis for helpful discussions and sharing *Arabidopsis* mutant lines and C. Campion and Prof. P. Simoneau for kindly providing the *Alternaria brassicicola* strain Abra43. Libraries and Miseq sequencing were performed by the GeT-Biopuces platform, INSA/TBI Genotoul Toulouse and member of IBISBA-FR (https://doi.org/10.15454/08BX-VJ91).

## AUTHOR CONTRIBUTIONS

CN, MZ, RP, AA, QB: experimental work, investigation, formal analysis, methodology. GM: supervision, editing. BD and TR: funding acquisition, project coordination, writing.

## Supplementary Table

**Supplementary Table S1.** Metabolomic dataset generated from ultra-high-performance liquid chromatography–high-resolution mass spectrometry (UHPLC-HRMS) analyses of Arabidopsis thaliana Col-0 and npr1 mutant extracts collected 10 days after inoculation with AgN23 spores or under mock conditions. Each detected feature is defined by its Cluster, Annotation, and Alignment ID (columns A–C). Retention time, ion m/z, and predicted XLogP are provided in columns D–F. A proposed metabolite name is given in column G, accompanied by an annotation score and annotation type (columns H–I). SMILES strings and additional structural identifiers appear in columns J–M. The proposed chemical structure for each compound is shown in column P. Log2 Fold Change (FC) comparing AgN23-treated versus mock-treated plants for both Col-0 and npr1 are presented in columns Q–T. Statistical significance for each ratio was assessed using a univariate Wilcoxon test comparing AgN23 and mock conditions within each genotype. ClassyFire natural product (NP) categories are reported in columns AC–AK. Raw abundance values for all detected metabolites are provided in columns AL–BO. The data is sorted by Retention Time, column E.

**Supplementary Table S2**. List of primers used in this study.

**Supplementary Table S3.** Differential abundance of ASVs in Arabidopsis thaliana Col-0 and npr1 mutant grown in potting soil under mock conditions or following AgN23 inoculation. Metabarcoding data were generated using an Illumina MiSeq platform with 250 bp paired-end sequencing targeting the V3–V4 region of the 16S rRNA gene, amplified with primers 341F (CCTACGGGNGGCWGCAG) and 805R (GACTACHVGGGTATCTAATCC). Raw FASTQ reads were processed using QIIME2 (v2021.4) with the DADA2 pipeline to obtain an amplicon sequence variant (ASV) table with taxonomic assignments. Differentially abundant ASVs were identified using the DESeq2 package in R (v3.4) via RStudio. Results are presented as log2 fold changes. Only ASVs with an adjusted p-value < 0.001 (Benjamini– Hochberg correction) are shown.

## REFERENCES

Agler, M. T., J. Ruhe, S. Kroll, C. Morhenn, S.-T. Kim, D. Weigel and E. M. Kemen (2016). "Microbial Hub Taxa Link Host and Abiotic Factors to Plant Microbiome Variation." PLOS Biology 14(1): e1002352.

Ahn, I.-P., S.-W. Lee and S.-C. Suh (2007). "Rhizobacteria-Induced Priming in Arabidopsis Is Dependent on Ethylene, Jasmonic Acid, and NPR1." Molecular Plant-Microbe Interactions® 20(7): 759–768.

Alfonso, E., E. Stahl, G. Glauser, E. Bellani, T. M. Raaymakers, G. Van den Ackerveken, J. Zeier and P. Reymond (2021). "Insect eggs trigger systemic acquired resistance against a fungal and an oomycete pathogen." New Phytologist 232(6): 2491–2505.

Banerjee, S. and M. G. A. van der Heijden (2023). "Soil microbiomes and one health." Nature Reviews Microbiology 21(1): 6–20.

Berg, G., D. Rybakova, D. Fischer, T. Cernava, M.-C. C. Vergès, T. Charles, X. Chen, L. Cocolin, K. Eversole, G. H. Corral, M. Kazou, L. Kinkel, L. Lange, N. Lima, A. Loy, J. A. Macklin, E. Maguin, T. Mauchline, R. McClure, B. Mitter, M. Ryan, I. Sarand, H. Smidt, B. Schelkle, H. Roume, G. S. Kiran, J. Selvin, R. S. C. d. Souza, L. van Overbeek, B. K. Singh, M. Wagner, A. Walsh, A. Sessitsch and M. Schloter (2020). "Microbiome definition re-visited: old concepts and new challenges." Microbiome 8(1): 103.

Bhattacharyya, A., D. Pablo Clint, V. Mavrodi Olga, S. Flynt Alex, M. Weller David, S. Thomashow Linda and V. Mavrodi Dmitri (2025). "Root exudates protect rhizosphere Pseudomonas from water stress." Applied and Environmental Microbiology 91(9): e00768–00725.

Brakhage, A. A. (2024). "Microbial Hub signaling compounds: natural products disproportionally shape microbiome composition and structure." microLife: uqae017.

Bulgarelli, D., M. Rott, K. Schlaeppi, E. Ver Loren van Themaat, N. Ahmadinejad, F. Assenza, P. Rauf, B. Huettel, R. Reinhardt, E. Schmelzer, J. Peplies, F. O. Gloeckner, R. Amann, T. Eickhorst and P. Schulze-Lefert (2012). "Revealing structure and assembly cues for Arabidopsis root-inhabiting bacterial microbiota." Nature 488(7409): 91–95.

Cao, H., S. A. Bowling, A. S. Gordon and X. Dong (1994). "Characterization of an Arabidopsis Mutant That Is Nonresponsive to Inducers of Systemic Acquired Resistance." The Plant Cell 6(11): 1583–1592.

Cao, H., J. Glazebrook, J. D. Clarke, S. Volko and X. Dong (1997). "The Arabidopsis NPR1 Gene That Controls Systemic Acquired Resistance Encodes a Novel Protein Containing Ankyrin Repeats." Cell 88(1): 57–63.

Carrión, V. J., J. Perez-Jaramillo, V. Cordovez, V. Tracanna, M. de Hollander, D. Ruiz-Buck, L. W. Mendes, W. F. J. van Ijcken, R. Gomez-Exposito, S. S. Elsayed, P. Mohanraju, A. Arifah, J. van der Oost, J. N. Paulson, R. Mendes, G. P. van Wezel, M. H. Medema and J. M. Raaijmakers (2019). "Pathogen-induced activation of disease-suppressive functions in the endophytic root microbiome." Science (New York, N.Y.) 366(6465): 606–612.

Cha, J. Y., S. Han, H. J. Hong, H. Cho, D. Kim, Y. Kwon, S. K. Kwon, M. Crüsemann, Y. Bok Lee, J. F. Kim, G. Giaever, C. Nislow, B. S. Moore, L. S. Thomashow, D. M. Weller and Y. S. Kwak (2016). "Microbial and biochemical basis of a Fusarium wilt-suppressive soil." ISME J 10(1): 119–129.

Chandrasekhar, V., K. Rajan, Sri Ram S. Kanakam, N. Sharma, V. Weißenborn, J. Schaub and C. Steinbeck (2025). "COCONUT 2.0: a comprehensive overhaul and curation of the collection of open natural products database." Nucleic Acids Research 53(D1): D634–D643.

Chaparro, J. M., D. V. Badri and J. M. Vivanco (2014). "Rhizosphere microbiome assemblage is affected by plant development." ISME J 8(4): 790–803.

Chapman, J. M. and G. K. Muday (2021). "Flavonols modulate lateral root emergence by scavenging reactive oxygen species in *Arabidopsis thaliana*." Journal of Biological Chemistry 296.

Chen, Y., J. Feng, P. Lü, D. Zhou, Y. Wei, T. Jing, Z. Zheng, W. Raza, D. Qi, M. Zhang, Y. Zhao, K. Li, W. Wang, X. Cheng and J. Xie (2026). "Streptomyces sesquiterpenes elicit 10-HCA secretion and recruit disease-suppressive microbiota to enhance banana Fusarium wilt resistance." Nature Communications.

Conn, V. M., A. R. Walker and C. M. Franco (2008). "Endophytic actinobacteria induce defense pathways in Arabidopsis thaliana." Mol Plant Microbe Interact 21(2): 208–218.

Diab, E., C. Du, W. Tigani, S. S. Elsayed and G. P. van Wezel (2026). "Plant Coumarins Modulate Natural Product Biosynthesis in a Streptomyces Root Endophyte." Journal of Natural Products.

Donald, L., A. Pipite, R. Subramani, J. Owen, R. A. Keyzers and T. Taufa (2022). "Streptomyces: Still the Biggest Producer of New Natural Secondary Metabolites, a Current Perspective." Microbiology Research 13(3): 418–465.

Durán, P., T. Thiergart, R. Garrido-Oter, M. Agler, E. Kemen, P. Schulze-Lefert and S. Hacquard (2018). "Microbial Interkingdom Interactions in Roots Promote Arabidopsis Survival." Cell 175(4): 973–983.e914.

Finkel, O. M., I. Salas-González, G. Castrillo, J. M. Conway, T. F. Law, P. J. P. L. Teixeira, E. D. Wilson, C. R. Fitzpatrick, C. D. Jones and J. L. Dangl (2020). "A single bacterial genus maintains root growth in a complex microbiome." Nature 587(7832): 103–108.

Fitzpatrick, C. R., J. Copeland, P. W. Wang, D. S. Guttman, P. M. Kotanen and M. T. J. Johnson (2018). "Assembly and ecological function of the root microbiome across angiosperm plant species." Proc Natl Acad Sci U S A 115(6): E1157–E1165.

Fitzpatrick, C. R., R. A. Smith, J. Hige, T. F. Law, D. Russ, O. E. Ajayi, A. A. Eida, P. Jacob, M. Jowers, N. Kumar, C. T. U. Lai, M. Anguita-Maeso, S. B. Peterson, C. Saha, T. Skelly, Q. Zhao, W. Zhou, S. R. Grant, J. D. Mougous, C. D. Jones and J. L. Dangl (2026). "*Streptomyces* enrichment in roots during drought is uncoupled from plant benefit and is driven by host suppression of iron uptake and immunity." bioRxiv: 2026.2003.2006.710171.

Fraisier-Vannier, O., J. Chervin, G. Cabanac, V. Puech-Pages, S. Fournier, V. Durand, A. Amiel, O. André, O. A. Benamar, B. Dumas, H. Tsugawa and G. Marti (2020). MS-CleanR: A feature-filtering approach to improve annotation rate in untargeted LC-MS based metabolomics, Bioinformatics.

Gao, M., C. Xiong, C. Gao, C. K. M. Tsui, M.-M. Wang, X. Zhou, A.-M. Zhang and L. Cai (2021). "Disease-induced changes in plant microbiome assembly and functional adaptation." Microbiome 9(1): 187.

Gayrard, D., C. Nicolle, M. Veyssière, K. Adam, Y. Martinez, C. Vandecasteele, M. Vidal, B. Dumas and T. Rey (2023). "Genome Sequence of the Streptomyces Strain AgN23 Revealed Expansion and Acquisition of Gene Repertoires Potentially Involved in Biocontrol Activity and Rhizosphere Colonization." PhytoFrontiers™: PHYTOFR-11-22-0131-R.

Getzke, F., T. Thiergart and S. Hacquard (2019). "Contribution of bacterial-fungal balance to plant and animal health." Curr Opin Microbiol 49: 66-72

Glazebrook, J. and F. M. Ausubel (1994). "Isolation of phytoalexin-deficient mutants of Arabidopsis thaliana and characterization of their interactions with bacterial pathogens." Proceedings of the National Academy of Sciences of the United States of America 91(19): 8955-8959.

Gu, S., Z. Wei, Z. Shao, V.-P. Friman, K. Cao, T. Yang, J. Kramer, X. Wang, M. Li, X. Mei, Y. Xu, Q. Shen, R. Kümmerli and A. Jousset (2020). "Competition for iron drives phytopathogen control by natural rhizosphere microbiomes." Nature Microbiology 5(8): 1002-1010.

Guzmán, P. and J. R. Ecker (1990). "Exploiting the triple response of Arabidopsis to identify ethylene-related mutants." The Plant Cell 2(6): 513-523.

Harbort, C. J., M. Hashimoto, H. Inoue, Y. Niu, R. Guan, A. D. Rombolà, S. Kopriva, M. J. E. E. E. Voges, E. S. Sattely, R. Garrido-Oter and P. Schulze-Lefert (2020). "Root-Secreted Coumarins and the Microbiota Interact to Improve Iron Nutrition in *Arabidopsis*." Cell Host & Microbe 28(6): 825-837.e826.

Huang, A. C., T. Jiang, Y.-X. Liu, Y.-C. Bai, J. Reed, B. Qu, A. Goossens, H.-W. Nützmann, Y. Bai and A. Osbourn (2019). "A specialized metabolic network selectively modulates Arabidopsis root microbiota." Science 364(6440): eaau6389.

Hull, A. K., R. Vij and J. L. Celenza (2000). "Arabidopsis cytochrome P450s that catalyze the first step of tryptophan-dependent indole-3-acetic acid biosynthesis." Proceedings of the National Academy of Sciences 97(5): 2379–2384.

Kai, K., M. Mizutani, N. Kawamura, R. Yamamoto, M. Tamai, H. Yamaguchi, K. Sakata and B.-i. Shimizu (2008). "Scopoletin is biosynthesized via ortho-hydroxylation of feruloyl CoA by a 2-oxoglutarate-dependent dioxygenase in Arabidopsis thaliana." The Plant Journal 55(6): 989–999.

Kempthorne, C. J., A. J. Nielsen, D. C. Wilson, J. McNulty, R. K. Cameron and D. K. Liscombe (2021). "Metabolite profiling reveals a role for intercellular dihydrocamalexic acid in the response of mature Arabidopsis thaliana to Pseudomonas syringae." Phytochemistry 187: 112747.

Kim, D. R., G. Cho, C. W. Jeon, D. M. Weller, L. S. Thomashow, T. C. Paulitz and Y. S. Kwak (2019). "A mutualistic interaction between Streptomyces bacteria, strawberry plants and pollinating bees." Nat Commun 10(1): 4802.

Koprivova, A., S. Schuck, R. P. Jacoby, I. Klinkhammer, B. Welter, L. Leson, A. Martyn, J. Nauen, N. Grabenhorst, J. F. Mandelkow, A. Zuccaro, J. Zeier and S. Kopriva (2019). "Root-specific camalexin biosynthesis controls the plant growth-promoting effects of multiple bacterial strains." Proceedings of the National Academy of Sciences 116(31): 15735–15744.

Koprivova, A., M. Schwier, V. Volz and S. Kopriva (2023). "Shoot-root interaction in control of camalexin exudation in Arabidopsis." J Exp Bot 74(8): 2667-2679.

Kost, C., K. R. Patil, J. Friedman, S. L. Garcia and M. Ralser (2023). "Metabolic exchanges are ubiquitous in natural microbial communities." Nature Microbiology 8(12): 2244–2252.

Krespach, M. K. C., M. C. Stroe, T. Netzker, M. Rosin, L. M. Zehner, A. J. Komor, J. M. Beilmann, T. Krüger, K. Scherlach, O. Kniemeyer, V. Schroeckh, C. Hertweck and A. A. Brakhage (2023). "Streptomyces polyketides mediate bacteria–fungi interactions across soil environments." Nature Microbiology 8(7): 1348–1361.

Lee, S.-M., H. G. Kong, G. C. Song and C.-M. Ryu (2021). "Disruption of Firmicutes and Actinobacteria abundance in tomato rhizosphere causes the incidence of bacterial wilt disease." The ISME Journal 15(1): 330-347.

Leon-Reyes, A., S. H. Spoel, E. S. De Lange, H. Abe, M. Kobayashi, S. Tsuda, F. F. Millenaar, R. A. M. Welschen, T. Ritsema and C. M. J. Pieterse (2009). "Ethylene Modulates the Role of NONEXPRESSOR OF PATHOGENESIS-RELATED GENES1 in Cross Talk between Salicylate and Jasmonate Signaling." Plant Physiology 149(4): 1797–1809.

Lewis, D. R., M. V. Ramirez, N. D. Miller, P. Vallabhaneni, W. K. Ray, R. F. Helm, B. S. J. Winkel and G. K. Muday (2011). "Auxin and Ethylene Induce Flavonol Accumulation through Distinct Transcriptional Networks." Plant Physiology 156(1): 144-164.

Lidbury, I. D. E. A., C. Borsetto, A. R. J. Murphy, A. Bottrill, A. M. E. Jones, G. D. Bending, J. P. Hammond, Y. Chen, E. M. H. Wellington and D. J. Scanlan (2020). "Niche-adaptation in plant-associated Bacteroidetes favours specialisation in organic phosphorus mineralisation." The ISME Journal: 1–16.

Ling, N., T. Wang and Y. Kuzyakov (2022). "Rhizosphere bacteriome structure and functions." Nature Communications 13(1): 836.

Liu, F., H. Jiang, S. Ye, W.-P. Chen, W. Liang, Y. Xu, B. Sun, J. Sun, Q. Wang, J. D. Cohen and C. Li (2010). "The Arabidopsis P450 protein CYP82C2 modulates jasmonate-induced root growth inhibition, defense gene expression and indole glucosinolate biosynthesis." Cell Research 20(5): 539–552.

Lu, H., C. Lu, S. Huang, W. Liu, L. Wang, C. Yang, E. Wang and L. Li (2025). "Rhizosphere microbes mitigate the shade avoidance responses in *Arabidopsis*." Cell Host & Microbe 33(6): 973-987.e974.

Lundberg, D. S., S. L. Lebeis, S. H. Paredes, S. Yourstone, J. Gehring, S. Malfatti, J. Tremblay, A. Engelbrektson, V. Kunin, T. G. d. Rio, R. C. Edgar, T. Eickhorst, R. E. Ley, P. Hugenholtz, S. G. Tringe and J. L. Dangl (2012). "Defining the core Arabidopsis thaliana root microbiome." Nature 488(7409): 86–90.

Lupini, A., F. Araniti, F. Sunseri and M. R. Abenavoli (2014). "Coumarin interacts with auxin polar transport to modify root system architecture in Arabidopsis thaliana." Plant Growth Regulation 74(1): 23–31.

Malka, S. K. and Y. Cheng (2017). "Possible Interactions between the Biosynthetic Pathways of Indole Glucosinolate and Auxin." Frontiers in Plant Science 8.

McLaughlin, S., K. Zhalnina, S. Kosina, T. R. Northen and J. Sasse (2023). "The core metabolome and root exudation dynamics of three phylogenetically distinct plant species." Nature Communications 14(1): 1649.

Nguyen, N. H., P. Trotel-Aziz, C. Clément, P. Jeandet, F. Baillieul and A. Aziz (2022). "Camalexin accumulation as a component of plant immunity during interactions with pathogens and beneficial microbes." Planta 255(6): 116.

Nicolle, C., D. Gayrard, A. Noël, M. Hortala, A. Amiel, S. Grat, A. Le Ru, G. Marti, J. L. Pernodet, S. Lautru, B. Dumas and T. Rey (2024). "Root-associated Streptomyces produce galbonolides to modulate plant immunity and promote rhizosphere colonization." ISME J.

O’Banion, B. S., P. Jones, A. A. Demetros, B. R. Kelley, L. H. Knoor, A. S. Wagner, J.-G. Chen, W. Muchero, T. B. Reynolds, D. Jacobson and S. L. Lebeis (2023). "Plant *myo*-inositol transport influences bacterial colonization phenotypes." Current Biology 33(15): 3111–3124.e3115.

Pastorczyk, M., A. Kosaka, M. Piślewska-Bednarek, G. López, H. Frerigmann, K. Kułak, E. Glawischnig, A. Molina, Y. Takano and P. Bednarek (2020). "The role of CYP71A12 monooxygenase in pathogen-triggered tryptophan metabolism and Arabidopsis immunity." New Phytologist 225(1): 400–412.

Pereira Francisco, V., S. Duthen, E. Crossay, A. l. Perez, S. Hennechart, M. Alignan, R. Valentin, F. Milone-Delacourt and G. Marti (2026). "MS-Net: Multi-Similarity-Based Network Annotation for Untargeted Metabolomics." Analytical Chemistry 98(23): 17116–17129.

Pieterse, C. M. J. (2025). "The Extended Plant Immune System." Molecular Plant-Microbe Interactions 38(6): 780-795.

Prout, J. N., A. Williams, A. Wanke, S. Schornack, J. Ton and K. J. Field (2024). "Mucoromycotina ‘fine root endophytes’: a new molecular model for plant–fungal mutualisms?" Trends in Plant Science 29(6): 650-661.

Pérez-Alonso, M.-M., P. Ortiz-García, J. Moya-Cuevas, T. Lehmann, B. Sánchez-Parra, R. G. Björk, S. Karim, M. R. Amirjani, H. Aronsson, M. D. Wilkinson and S. Pollmann (2021). "Endogenous indole-3-acetamide levels contribute to the crosstalk between auxin and abscisic acid, and trigger plant stress responses in Arabidopsis." Journal of Experimental Botany 72(2): 459-475.

Rajniak, J., B. Barco, N. K. Clay and E. S. Sattely (2015). "A new cyanogenic metabolite in Arabidopsis required for inducible pathogen defence." Nature 525(7569): 376-379.

Russ, D., C. R. Fitzpatrick, P. J. P. L. Teixeira and J. L. Dangl (2023). "Deep discovery informs difficult deployment in plant microbiome science." Cell 186(21): 4496-4513.

Růžička, K., K. Ljung, S. Vanneste, R. Podhorská, T. Beeckman, J. i. Friml and E. Benková (2007). "Ethylene Regulates Root Growth through Effects on Auxin Biosynthesis and Transport-Dependent Auxin Distribution." The Plant Cell 19(7): 2197-2212.

Sellam, A., A. Dongo, T. Guillemette, P. Hudhomme and P. Simoneau (2007). "Transcriptional responses to exposure to the brassicaceous defence metabolites camalexin and allyl-isothiocyanate in the necrotrophic fungus Alternaria brassicicola." Molecular Plant Pathology 8(2): 195-208.

Sharma, I., S. Kashyap and N. Agarwala (2023). "Biotic stress-induced changes in root exudation confer plant stress tolerance by altering rhizospheric microbial community." Frontiers in Plant Science 14.

Stassen, M. J. J., S.-H. Hsu, C. M. J. Pieterse and I. A. Stringlis (2021). "Coumarin Communication Along the Microbiome-Root-Shoot Axis." Trends in Plant Science 26(2): 169–183.

Staswick, P. E., W. Su and S. H. Howell (1992). "Methyl jasmonate inhibition of root growth and induction of a leaf protein are decreased in an Arabidopsis thaliana mutant." Proceedings of the National Academy of Sciences 89(15): 6837-6840.

Stringlis, I. A., R. de Jonge and C. M. J. Pieterse (2019). "The Age of Coumarins in Plant-Microbe Interactions." Plant Cell Physiol 60(7): 1405–1419.

Sánchez-Parra, B., M.-M. Pérez-Alonso, P. Ortiz-García, J. Moya-Cuevas, M. Hentrich and S. Pollmann (2021) "Accumulation of the Auxin Precursor Indole-3-Acetamide Curtails Growth through the Repression of Ribosome-Biogenesis and Development-Related Transcriptional Networks." International Journal of Molecular Sciences 22 DOI: 10.3390/ijms22042040.

Trivedi, P., J. E. Leach, S. G. Tringe, T. Sa and B. K. Singh (2020). "Plant–microbiome interactions: from community assembly to plant health." Nature Reviews Microbiology 18(11): 607-621.

Tsai, H.-H., Y. Tang, L. Jiang, X. Xu, V. Dénervaud Tendon, J. Pang, Y. Jia, K. Wippel, J. Vacheron, C. Keel, T. G. Andersen, N. Geldner and F. Zhou "Localized glutamine leakage drives the spatial structure of root microbial colonization." Science 390(6768): eadu4235.

Tsugawa, H., T. Cajka, T. Kind, Y. Ma, B. Higgins, K. Ikeda, M. Kanazawa, J. VanderGheynst, O. Fiehn and M. Arita (2015). "MS-DIAL: data-independent MS/MS deconvolution for comprehensive metabolome analysis." Nature Methods 12(6): 523–526.

Tsugawa, H., T. Kind, R. Nakabayashi, D. Yukihira, W. Tanaka, T. Cajka, K. Saito, O. Fiehn and M. Arita (2016). "Hydrogen Rearrangement Rules: Computational MS/MS Fragmentation and Structure Elucidation Using MS-FINDER Software." Analytical chemistry 88(16): 7946-7958.

Van der Ent, S., B. W. Verhagen, R. Van Doorn, D. Bakker, M. G. Verlaan, M. J. Pel, R. G. Joosten, M. C. Proveniers, L. C. Van Loon, J. Ton and C. M. Pieterse (2008). "MYB72 is required in early signaling steps of rhizobacteria-induced systemic resistance in Arabidopsis." Plant Physiol 146(3): 1293–1304.

Vergnes, M. S., M. D. Gayrard, M. M. Veyssiere, M. J. Toulotte, M. Y. Martinez, D. V. Dumont, D. O. Bouchez, D. T. Rey and D. B. Dumas (2020). "Phyllosphere colonisation by a soil Streptomyces sp. promotes plant defense responses against fungal infection." 43.

Xiong, X., J. Zeng, Q. Ning, H. Liu, Z. Bu, X. Zhang, J. Zeng, R. Zhuo, K. Cui, Z. Qin, Y. Gao, X. Liu and Y. Zhu (2024). "Ferroptosis induction in host rice by endophyte OsiSh-2 is necessary for mutualism and disease resistance in symbiosis." Nature Communications 15(1): 5012.

Xu, L., D. Naylor, Z. Dong, T. Simmons, G. Pierroz, K. K. Hixson, Y.-M. Kim, E. M. Zink, K. M. Engbrecht, Y. Wang, C. Gao, S. DeGraaf, M. A. Madera, J. A. Sievert, J. Hollingsworth, D. Birdseye, H. V. Scheller, R. Hutmacher, J. Dahlberg, C. Jansson, J. W. Taylor, P. G. Lemaux and D. Coleman-Derr (2018). "Drought delays development of the sorghum root microbiome and enriches for monoderm bacteria." Proceedings of the National Academy of Sciences 115(18): E4284–E4293.

Yang, Z., Y. Qiao, N. C. Konakalla, E. Strøbech, P. Harris, G. Peschel, M. Agler-Rosenbaum, T. Weber, E. Andreasson and L. Ding (2023). "Streptomyces alleviate abiotic stress in plant by producing pteridic acids." Nature Communications 14(1): 7398.

Yu, K., H. Liu, W. Zhong and I. A. Stringlis (2022). Microbiome-assisted Agriculture. Biocontrol of Plant Disease, John Wiley & Sons, Ltd: 217–253.

Yu, K., Y. Liu, R. Tichelaar, N. Savant, E. Lagendijk, S. J. L. van Kuijk, I. A. Stringlis, A. J. H. van Dijken, C. M. J. Pieterse, P. A. H. M. Bakker, C. H. Haney and R. L. Berendsen (2019). "Rhizosphere-Associated Pseudomonas Suppress Local Root Immune Responses by Gluconic Acid-Mediated Lowering of Environmental pH." Curr Biol 29(22): 3913–3920.e3914.

Yuan, J., J. Zhao, T. Wen, M. Zhao, R. Li, P. Goossens, Q. Huang, Y. Bai, J. M. Vivanco, G. A. Kowalchuk, R. L. Berendsen and Q. Shen (2018). "Root exudates drive the soil-borne legacy of aboveground pathogen infection." Microbiome 6(1): 156.

Zhao, Q., S. Bertolli, Y.-J. Park, Y. Tan, K. J. Cutler, P. Srinivas, K. L. Asfahl, C. Fonesca-García, L. A. Gallagher, Y. Li, Y. Wang, D. Coleman-Derr, F. DiMaio, D. Zhang, S. B. Peterson, D. Veesler and J. D. Mougous (2024). "Streptomyces umbrella toxin particles block hyphal growth of competing species." Nature 629(8010): 165–173.

Zhao, Y., A. K. Hull, N. R. Gupta, K. A. Goss, J. Alonso, J. R. Ecker, J. Normanly, J. Chory and J. L. Celenza (2002). "Trp-dependent auxin biosynthesis in Arabidopsis: involvement of cytochrome P450s CYP79B2 and CYP79B3." Genes & Development 16(23): 3100–3112.

Zhou, N., T. L. Tootle and J. Glazebrook (1999). "Arabidopsis PAD3, a Gene Required for Camalexin Biosynthesis, Encodes a Putative Cytochrome P450 Monooxygenase." The Plant Cell 11(12): 2419–2428.

